# Mass allelic exchange: enabling sexual genetics in *Escherichia coli*

**DOI:** 10.1101/2020.11.30.405282

**Authors:** Varnica Khetrapal, Liyana Ow Yong, Swaine L. Chen

## Abstract

Despite dramatic advances in genomics, connecting genotypes to phenotypes is still challenging. Sexual genetics combined with linkage analysis is a powerful solution to this problem but generally unavailable in bacteria. We build upon a strong negative selection system to invent Mass Allelic Exchange (MAE), which enables hybridization of arbitrary (including pathogenic) strains of *E. coli*. MAE reimplements the natural phenomenon of random crossovers, enabling classical linkage analysis. We demonstrate the utility of MAE with virulence-related gain-of-function screens, discovering that transfer of a single operon from a uropathogenic strain is sufficient for enabling a commensal *E. coli* to form large intracellular bacterial collections within bladder epithelial cells. MAE thus enables assaying natural allelic variation in *E. coli* (and potentially other bacteria), complementing existing loss-of-function genomic techniques.

**One Sentence Summary:** We create F1 hybrids of *E. coli* using MAE, bringing the power of linkage analysis to bear on phenotypic diversity (including virulence)

## Introduction

Genetics is one of the foundational engines upon which modern biological knowledge has been built. Early seminal discoveries and innovations, including the systematic analysis of heritability experiments by Mendel (*1*), created the context in which subsequent discoveries of the structure of DNA, the central dogma, and the first DNA sequencing strategies became biological canon (*2–5*). Genetics is the field that is worked by evolution, the concept without which nothing makes sense (*6*).

Our ability to harness experimental genetics has had two major technological drivers: natural processes and engineered systems/applications. Examples of the former include Mendel’s early experiments with crossing pea plants, directed breeding to generate and optimize numerous domesticated plants and animals, and selection of spontaneous mutants of bacteria with antibiotics or phage. The latter category, where natural processes are harnessed for sometimes drastically different purposes, has accelerated in step with our understanding of biology. Examples of such engineered systems include a wide array of engineered plasmid-based technology (*7*), numerous transposon-mediated strategies (ranging from P element-, sleeping beauty-, and piggybac-mediated mutagenesis in eukaryotic systems to Tn-seq (transposon-sequencing) in bacteria) (*8–12*), Type IV secretion-mediated transformation of plants (*13*), and the recent and dramatic explosion of CRISPR-Cas-based applications (for both mutagenesis, regulation, and even facilitating DNA sequencing) (*14–16*).

Of note, many of the advances in genetic technology from the last several decades have been derived from bacterial systems. To focus on genetics in bacteria, we in principle have a full suite of capabilities to both sequence and synthesize complete bacterial genomes (*17, 18*). Unfortunately, our technical capabilities outstrip our knowledge of what these sequences do.

Even in *Escherichia coli*, the flagship workhorse of molecular genetics, biochemistry, and synthetic biology, it is estimated that we do not understand the function of up to 35% of its genes (*19*). Therefore, each additional technology, particularly those enabling high throughput characterization, has driven deeper insights and a broader understanding of biology. Recently, Tn-seq has been a revolutionary tool that has modernized traditional genetic screens, simultaneously providing speed, completeness, and quantitation for identifying the genetic contributors to a given phenotype, ranging from simple growth under certain conditions to the ability to cause disease (*10*).

Unfortunately, except in rare cases, traditional screens and Tn-seq provide only one type of information we need to understand (and subsequently engineer) genetic pathways. These both largely explore the space of loss-of-function mutations. They also are largely blind to phenotypes driven by essential genes. Modifications to Tn-seq (with outward facing promoters) (*20, 21*) or use of plasmid-based overexpression libraries can partially enable gain-of-function screens, though these raise additional issues such as artificial expression levels and regulation. Other alternatives available recently are enabled by advanced techniques such as CAGE, MAGE, REGRES, REXER, and GENESIS (*22–25*); the power of these has been recently demonstrated with complete recoding to remove 3 codons throughout the *E. coli* genome (*26*). However, these technologies variously suffer from being heavily dependent on well-characterized (and “well-behaved”) cloning strains of *E. coli*; being unable to capture large insertions (such as pathogenicity islands, which are fundamentally important for bacterial evolution and genetics); requiring residual markers or recombination scars, limiting the potential for iteration; or not having been implemented at a genome-wide scale, which often uncovers additional technical limitations.

All of these issues could be solved by a traditional approach of hybridizing two bacterial strains that differ in phenotypes; this is the same conceptual framework that drove the initial directed hybridization experiments of Mendel. We previously demonstrated the power of bacterial sexual genetics using the naturally competent bacterium *Campylobacter jejuni* (*27*). By creating a library of sexual hybrids (akin to F1 progeny) between an abortifacient and a non-abortifacient strain, we deduced from a single animal infection experiment that mutations in one gene, *porA*, were responsible for the ability of *C. jejuni* to cause abortion. We were able to then experimentally demonstrate that specific allelic variants of *porA* were both necessary and sufficient for this disease phenotype (*27*).

Unfortunately, most bacteria are not naturally competent. We previously have leveraged dedicated toxin genes to create a modular and stringent negative selection system for use in *E. coli* and other Enterobacteriaceae; this system can be 1000× more stringent in its selection strength than other negative selection systems that can be used in unmodified hosts (*28*). Driven by the “selective headroom” provided by this advance in stringency, we have now combined this with transposon technology and Hfr (high frequency recombination) conjugation to create, for the first time, an engineered process to create complex libraries of unmarked sexual hybrids of *E. coli*. We term our new method Mass Allelic Exchange (MAE).

## Results and Discussion

A conceptual schematic of MAE is shown in **Fig. 1A**. We designate one *E. coli* strain as the donor and the other as the recipient. Both are subjected to transposon mutagenesis with a customized transposon which carries a stringent negative selection cassette. The donor transposon additionally has an outward facing *oriT* sequence to initiate chromosomal transfer adjacent to (and not including) the custom transposon. For tracking efficiency, the donor and recipient transposons also differ in their positively selectable antibiotic markers and a fluorescent protein (mCherry for donors, a plasmid-based GFP for recipients). Finally, the donor is transformed with a helper plasmid for conjugation, then a standard conjugation is performed. Selection of the mixture on restrictive conditions for the negative selection cassette results in survival only of hybrid transconjugant strains; the donor negative selection cassette is not transferred due to the position and orientation of the *oriT*, while the recipient negative selection cassette is removed by recombination with incoming donor DNA. Of note, we had initially considered using transformation of purified genomic DNA (as was done with *C. jejuni*) or generalized transduction as methods to exchange DNA between strains; using these, the donor strain would not need to be modified. However, transformation, even in *E. coli*, was found to be too inefficient by several orders of magnitude, while transduction required strain-specific phages that are not generally available for many *E. coli* strains, especially clinical isolates (*29*).

**Fig. 1.**
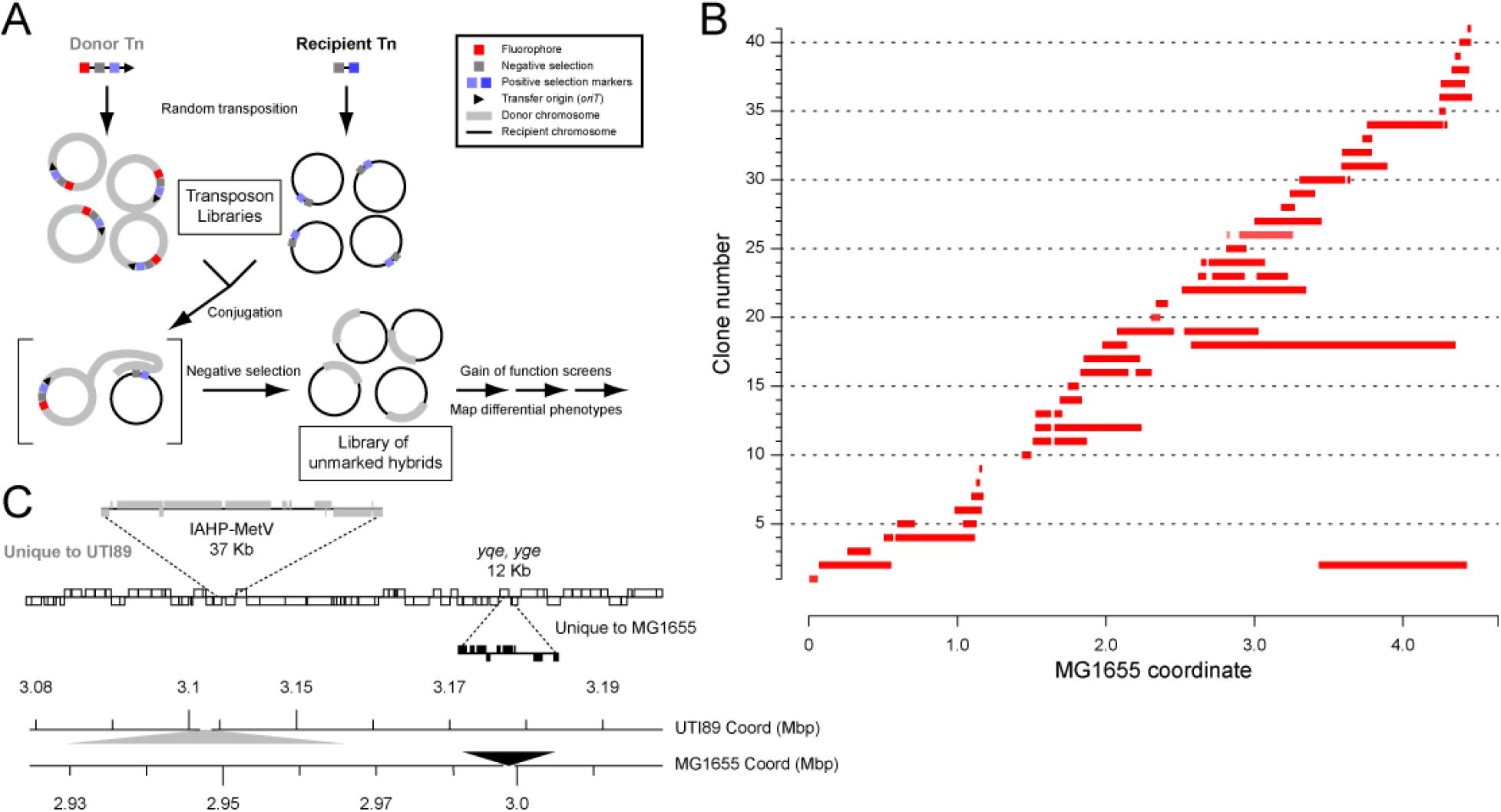
Overview of Mass Allelic Exchange. (**A**) Schematic of the MAE protocol. See text for further details. Each circle represents a single bacterial chromosome within a cell (for clarity, cell membranes are not drawn). Genetic markers are represented by colored squares, according to the legend at the top right. Experimental manipulations are indicated by the text adjacent to arrows. Text boxes indicate libraries of clones. Brackets enclose a representative single donor and recipient conjugation in the bottom left section. (**B**) Transferred SNPs for the SLC-H1 library of hybrids. Individual transconjugants analyzed using the MG1655 genome as a reference. Genomic coordinate is plotted on the x-axis. Individual clones are plotted at different locations on the y-axis. Blocks of consecutive single nucleotide polymorphisms (SNPs) relative to the MG1655 genome sequence are represented by red bars. (**C**) One transconjugant was assembled and compared to the UTI89 and MG1655 genome reference sequences. A comparison of the annotations of approximately 100 Kb of homologous regions of the UTI89 and MG1655 chromosomes is shown. Annotated genes are indicated by rectangles in the upper graph; shared genes are indicated by open rectangles, with coding strand indicated by position of the rectangle above (positive) or below (negative) the central line. Genes found only in UTI89 are indicated by gray rectangles and offset upwards. Genes found only in MG1655 are indicated by black rectangles and offset downwards. The genomic coordinates for each reference are shown on dual scale at the bottom. Filled gray (for UTI89) and black (for MG1655) triangles indicate the location and length of large DNA insertions relative to the other strain. In the ~100 Kb region depicted, the transconjugant matches the UTI89 sequence, with assembled sequence matching the IAHP-MetV pathogenicity island but not the *yqe, yge* island.

The key innovation of MAE is the use of a stringent negative selection cassette, which has enough selective power to nearly quantitatively eliminate the numerically dominant, unhybridized donor and recipient cells; in a single step, we can thereby generate unmarked hybrids, which sets MAE apart from all previous technologies. Therefore, a practical issue limiting MAE was an apparent loss of negative selection stringency when using transposon libraries (10^-4^) instead of targeted insertions (10^-7^ - 10^-8^) (**Fig. S1A**). Growth under restrictive conditions of strains carrying the negative selection cassette can occur for multiple reasons, such as organic mutation of the selection cassette, inactivation via insertion sequences or other mobile elements, or spontaneous deletion. Certain regions of the chromosome are known to have a high effective spontaneous deletion rate, such as prophages, pathogenicity islands, and plasmids (*30, 31*) **Fig. S1A**). We therefore solved this issue by screening individual colonies for their frequency of growth on restrictive conditions. Aggregation of the “stable” donor and recipient colonies (with an inactivation frequency of <10^-6^) then resulted in a donor and recipient library with an overall negative selection stringency similar to that previously reported (10^-7^) (**Supplemental text 1**). Of note, this process secondarily resulted in a novel, rapid screen for inherently unstable regions of the *E. coli* chromosome; as expected, prophages, PAIs, and the plasmid are lost at relatively high frequency. Ribosomal RNA loci, which are known to undergo intra-chromosomal gene conversion, were also found. In addition, we discovered previously unknown unstable regions that may provide new insights into mechanisms of chromosome plasticity, maintenance, and evolution (**Supplemental text 1, Fig. S1B**).

In a single hybridization experiment, we were able to generate thousands of hybrid transconjugants (based on blocks of SNPs transferred, **Fig. S2A**). Based on resistance phenotypes, fluorescent markers, and whole genome sequencing, we estimate that this pooled MAE library is composed of >98% true hybrid transconjugants. We saw a bias towards DNA exchange near the origin of replication (**Fig. S2A**); this suggested that DNA replication, and the attendant differences in copy number between the origin and the terminus, was responsible (*32*). Indeed, we saw typical (3-7x) biases in the density of transposon insertions around the origin in the source donor and recipient libraries (**Fig. S2B** and **Fig. S2C**); conjugation and sequencing then incur additional bias amplifying origin-proximal transposon insertions and the resulting hybrids (**Fig. S2A**). A quantitative analysis of the bias across both source libraries and the output transconjugant library indicated that the origin copy number effect might be sufficient to account for all of the observed bias (**Supplemental text 2**). To verify that terminus-proximal crossovers were not further selected against, we performed directed transfers using individual donor and recipient strains, which showed no difference in efficiency (transconjugants per recipient) when the transfer was origin-proximal or terminus-proximal (**Fig. S2D** and **Fig. S2E**). In other words, the actual hybridization step in MAE introduced no additional bias. Therefore, control of growth rate (to limit multifork replication) and/or selection of uniformly (or terminus-biased) clones for the source libraries would effectively eliminate this bias. Alternatively, clone selection could be done after hybridization, particularly to create arrayed libraries useful for many genetic screens. We created the first such arrayed library of UTI89-MG1655 hybrids (SLC-H1), in which 88% of the UTI89 chromosome is represented in at least one clone (the largest missing block being 262 Kbp (5.6% of the genome)) (**Fig. 1B**).

All mutation types were accessible to transfer by MAE. Whole genome sequencing demonstrated continuous tracts of donor SNPs being transferred to the homologous region of the recipient chromosome. These were full replacements of segments of the chromosome, ranging from 16-1780Kb (median 298kb) (**Fig. 1B, Fig. S2A,** and **Supplemental text 3**), and they transferred all expected gene insertions and deletions as well as pathogenicity islands up to 80 kb (**Fig. 1C**). Most (273/372, 73.4%) hybrids had only one single block of DNA transferred, with additional separate blocks progressively more rare (**Fig. 1B, Fig. S2A,** and **Fig. S3**).

One of the key advantages of allelic exchange methods over the popular Tn-seq or other random mutagenesis methods is the ability to select for phenotypes that differ between strains, particularly in a gain-of-function strategy. We used pooled MAE libraries to perform two such screens. The donor strain, UTI89, survives in human serum, resists P1 phage infection, and hemolyzes sheep blood, while the recipient strain, MG1655, lacks all of these phenotypes. The first two phenotypes are known to be due to an insertion sequence disrupting the MG1655 *wbbL* gene, which abolishes LPS O-antigen synthesis (*33*). Interestingly, the UTI89 *rfb* locus is organized differently from the corresponding MG1655 locus; restoration of O-antigen in MG1655 should therefore require replacement of the entire 10kb locus (**Fig. 2A**). We first screened the pooled library for serum resistance, isolating 60 clones (**Fig. 2B, Fig. S4A**). Whole genome sequencing of 22 of these clones showed that only 3 independent transconjugants were isolated; in these, while the majority of the chromosome was indeed MG1655, 15-50kB sections of the UTI89 genome encompassing the *rfb* locus were now integrated into the homologous location of MG1655 (**Fig. 2C**). As further confirmation, these clones had also gained resistance to P1 phage, as expected (**Fig. S4B**). Finally, we verified separately that a targeted transfer of the entire *rfb* locus from UTI89 to MG1655 indeed is sufficient to confer resistance to serum and P1 phage infection (**Fig. S5**).

**Fig. 2.**
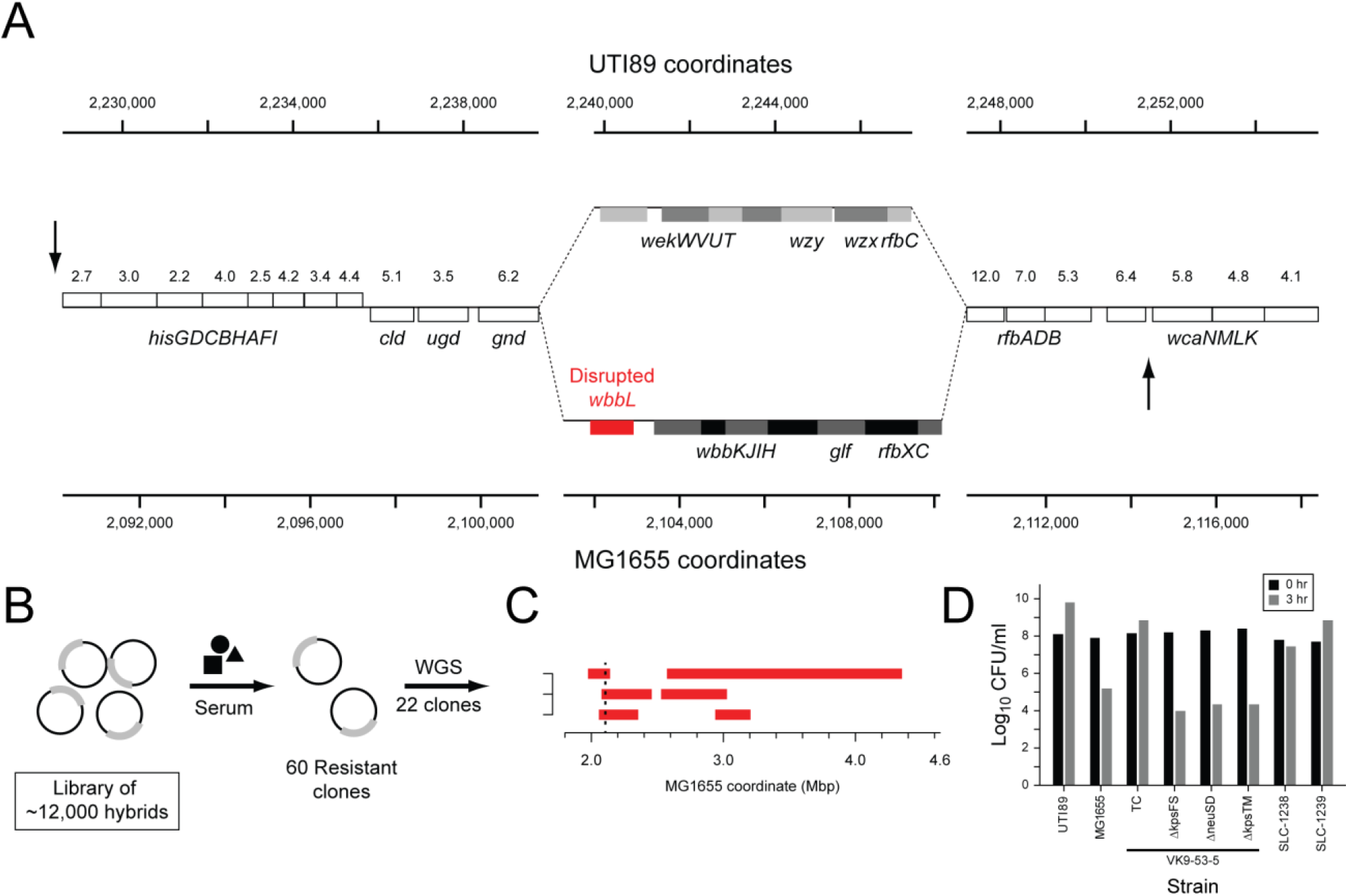
*In vitro* screening of MAE Libraries. (**A**) *rfb* locus comparison. Genome coordinates for UTI89 and MG1655 are shown on the axes at the top and bottom, respectively. Genes indicated by open boxes are nearly identical in sequence between UTI89 and MG1655; the percent nucleotide divergence is indicated by the numbers above each gene. Genes in the center section (gray boxes and red box) are not obviously homologous, with no sequence above 70% nucleotide identity in the other genome. Arrows indicate locations of donor (top arrow pointing down) and recipient (bottom arrow pointing up) insertions for targeted transfer. (**B**) Schematic of the serum and phage resistance screens. Each black circle represents a single bacterial chromosome within a cell (for clarity, cell membranes are not drawn). Experimental manipulations are indicated by the text adjacent to arrows. Thicker gray lines represent donor DNA sequence recombined into the recipient chromosome. (**C**) 22 serum resistant clones were sequenced, which were represented by three independent clones. The transferred regions for each clone are shown in red, based on the MG1655 coordinates (x-axis). The two vertical black dotted lines (very closely spaced) indicate the region of the *rfb* locus from the disrupted *wbbL* gene to the *rfbXC* genes. (**D**) Quantification of phage resistance. Each strain was titered before (0 hr, black) and 3 hours after (3 hr, gray) mixing with P1 phage; the logarithm of the titers are plotted on the x-axis. UTI89 is a resistant control (resistant to P1); MG1655 is a sensitive control. Strain VK-9-53-5 is a phage resistant transconjugant; this strain (TC) and derivatives with the indicated genes knocked out were also tested. Finally, SLC-1238 and SLC-1239 are two transconjugant strains derived from a directed transfer near the *kps* locus.

Attesting to the ability of gain of function screens to provide complementary information, MAE screens provided new insight into these well studied phenotypes. In control experiments to prepare for an intracellular infection screen (see below), we isolated phage-resistant clones (**Fig. 2D**) in which the *rfb* locus had not been transferred; instead, a region of the chromosome 1 Mbp away, encompassing a UTI89-specific 30 Kb island containing the *kps* and *neu* operons encoding a capsule biosynthetic pathway, had been transferred (VK9-53-5). A subsequent targeted transfer of this *kps*-containing locus (strains SLC-1238 and SLC-1239) showed that it does indeed confer an intermediate level of phage resistance (i.e. resistance only when infected with <10^10^ pfu/ml of P1 phage) (**Fig. 2D**). To verify that this was due to the capsule genes, we made several targeted deletions within the *kps* operon, all of which have been previously shown to abolish capsule production; these led to high phage sensitivity in the originally isolated resistant transconjugant (**Fig. 2D**). This is the first demonstration that modification of the capsule, in the absence of restoration of O-antigen, can confer partial resistance to P1 phage in MG1655.

Finally, we used MAE to explore a suspected complex phenotype: intracellular infection of cultured human epithelial cells. UTI89 is a uropathogenic strain; it robustly infects the bladder of mice and forms intracellular bacterial communities (IBC) during acute phases of infection (*34*). IBC formation imposes a severe population bottleneck (> 10^4^ reduction) on the inoculated UTI89 (*35*). This bottleneck precludes many genetic techniques to probe the mechanism of intracellular infection; therefore, several attempts have been made to create a similar infection system using *in vitro* cultured cells (*36, 37*). One of these utilizes saponin treatment of infected bladder epithelial cells (5637). In this system, UTI89 forms large intracellular aggregates of bacteria that are morphologically reminiscent of IBCs; these *in vitro* aggregates are referred to as pods. MG1655, in contrast, does not infect mouse bladders and also does not form pods *in vitro* (*36*). Therefore, infection of 5637 cells and treatment with saponin can be used as a gain-of-function screen for pod formation (**Fig. 3A**). We indeed found that pods were formed when we infected 5637 cells with our MAE library. We isolated 5 individual cells from two different experiments containing pods from the hybrid-infected cells; whole genome sequencing revealed a common 314 Kb region of the UTI89 chromosome that had been transferred in all clones (**Fig. 3B**). This region contained 6,148 SNPs, 35 genes within 4 operons present in UTI89 but not MG1655, another island of 34 translocated genes (i.e. they are also present in MG1655 but at a different chromosomal location), and 3 genes within 1 operon present in MG1655 but not in UTI89 (**Fig. 3C**). To further map this 314 Kb region, we screened another 102 individual transconjugants using PCRs for 6 genes distributed across this 314 Kb region that were present only in UTI89 or MG1655. 18 transconjugants were found that appeared to have transferred at least some of the 314 Kb locus (with 6 having transferred all) (**Fig. 3B**, pink and black bars). These 18 transconjugants were sequenced and also tested for pod formation; only 36 Kb was now shared among all 9 pod-forming clones. This 36 Kb pod-associated locus contained a total of 20 genes shared between UTI89 and MG1655, with average nucleotide identity of 96.9%; and 2 operons (encoding a PTS system and the *chu* heme acquisition system) that were present only in UTI89.

**Fig. 3.**
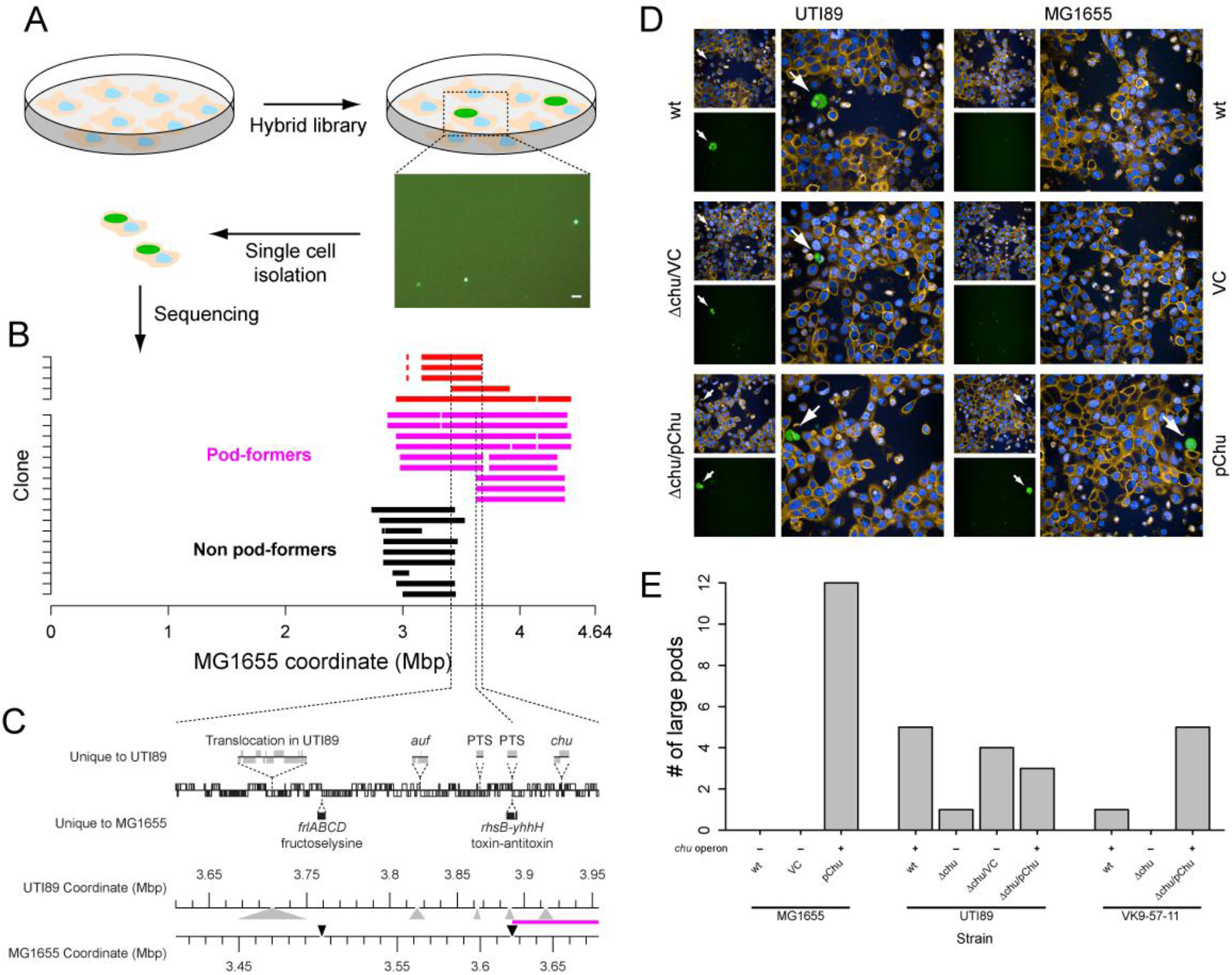
A gain-of-function screen for *in vitro* pod formation. (**A**) Schematic of the screen. Orange shapes represent individual bladder epithelial cells, light blue represents nuclei, and green represents aggregates of fluorescent *E. coli*. A representative image of green fluorescence (*E. coli*) after infection and saponin treatment is shown. Scale bar is 100 μm. (**B**) Whole genome sequencing of 5 pod-forming transconjugants and 18 additional transconjugants (see text for details) (y-axis). The transferred regions for each clone are shown in red or magenta (pod-forming transconjugants) or black (non-pod-forming transconjugants), based on the MG1655 coordinates (x-axis). The left-most and right-most vertical dotted lines indicates the common region of UTI89 transferred to the clones depicted in red. The middle vertical dotted line delimits the left border of the common region of UTI89 transferred to all pod-forming transconjugants. (**C**) Genetic map of the common overlapping region for the 5 pod-forming transconjugants. Notations are as in Figure 1C. Magenta line between the UTI89 and MG1655 coordinate axes at the bottom denotes the common overlapping region of all pod-forming clones. (**D**) Representative images of the pod-forming assay. Each strain is represented by three images; the smaller images on the left are the DAPI (blue) and wheat germ agglutinin (orange) channels (top) and GFP (green) (bottom). The larger image is a merge of all channels. White arrows indicate identified large pods. (**E**) The number of large pods (> 10,000 pixels) formed by each strain (total results for n=8 independent experiments). The parental strain is shown on the x-axis, along with the genotype. VC, vector control; pChu, plasmid containing the *chu* operon. The presence/absence of a complete *chu* operon is indicated by +/- above the strain genotypes.

We first tested the hypothesis that the UTI89-specific operons were sufficient for conferring pod formation to MG1655. We transformed plasmids carrying each of these operons into MG1655; only the strain carrying the UTI89 *chu* operon was able to form pods (**Fig. 3D, Movie S1, Fig. S6B**, and data not shown). In addition, deletion of the *chu* operon (but not the *auf* and the *PTS* operon; data not shown) from the transconjugant clones eliminated large pods (**Fig. 3E, Fig. 6SA**). Consistent with previous reports examining IBCs in mouse bladders (*38*), deletion of the *chu* operon in UTI89 led to smaller pods formed by UTI89, attributable to delayed kinetics of pod growth in the epithelial cells **(Fig. 6SC, Movie S2)**. We therefore conclude that the *chu* operon is sufficient to confer pod formation to MG1655.

Notably, MAE libraries coupled with this gain-of-function screening strategy were critical to make this discovery of the sufficiency of the *chu* operon. Previous gene expression- and loss-of-function-based strategies had identified this locus playing a quantitative but not necessary role in intracellular infection in UTI89. Perhaps because many other loci (prominent among which is the *fim* operon) are indeed necessary for intracellular infection (based on loss-of-function experiments), no further studies have examined the role of the *chu* operon in intracellular infection. The additional genetic information about the sufficiency of the *chu* operon therefore provides a key complementary insight into the mechanism of intracellular infection, which may focus future research efforts.

In summary, we have for the first time created intentional, unmarked whole genome hybrids, akin to F1 progeny, between two distinct strains of *E. coli*, in a process we term MAE. The core enabling technology for MAE is the negative selection system, which provides high selective headroom and is usable in all tested *E. coli* strains, as well as in Salmonella, *Serratia marcescens, Shigella flexneri, Enterobacter cloacae*, (*39*), *Providencia stuartii* (*40*), and Klebsiella (Y.H. Gan, personal communication) strains, including clinical isolates. This opens the possibility of applying the power of traditional sexual genetics to understanding complex phenotypes, such as infection, in the non-naturally competent Enterobacteriaceae. As with traditional sexual genetics, MAE provides access to all types of genetic variation between strains, from single nucleotide polymorphisms to large (>100 Kb) chromosomal insertions and deletions. Furthermore, MAE tests existing wild type sequences, as opposed artificial deletions or disruptions, and there are no residual genetic scars or markers in the resulting hybrid strains. Gain-of-function experiments are a natural application area for MAE, which complements other techniques that enable loss-of-function experiments. In addition to providing insights into previously inaccessible (or difficult-to-access) complex phenotypes, MAE also provides an additional technology to support the directed evolution of bacterial strains, enabling strategies akin to dog breeding for making customized chassis for biotechnology or synthetic biology applications.

## Acknowledgments

Whole genome sequencing was done at Genome Institute of Singapore, A*STAR. We thank Jane Vin Chan and Judice Koh at the High Throughput Phenomics, Genome Institute of Singapore for their support in microscopic and quantitative analysis of saponin facilitated pods. We thank members of the Chen lab, Kimberly Kline, Shu Sin Chng, and Emanuel Hanski for helpful discussions and suggestions on the manuscript. **Funding:** National Research Foundation, Prime Minister’s Office, Singapore under its NRF Research Fellowship Scheme [NRF-RF2010-10 to SLC]; National Medical Research Council, Ministry of Health, Singapore [NMRC/CIRG/1358/2013 and NMRC/OFIRG/0009/2016 to SLC]; Genome Institute of Singapore (GIS)/Agency for Science, Technology and Research (A*STAR). Funding for open access charge: National Medical Research Council, Ministry of Health, Singapore [NMRC/OFIRG/0009/2016].

## Author contributions

VK and SLC conceptualised the study. VK and LOY carried out the experiments. VK, LOY, and SLC analyzed data. VK and SLC wrote the manuscript.

## Data and materials availability

All data is available in the main text or the supplementary materials.

## Supplementary Materials

Materials and Methods

Supplementary Text 1 to 6

Figs. S1 to S6

Tables S1 to S3

Captions for Movies S1 to S2

