## Supplemental Materials for "Mass allelic exchange: enabling sexual genetics in *Escherichia coli*"

**This PDF file includes:**

Materials and Methods

Supplementary Text 1 to 6

Figs. S1 to S6

Tables S1 to S3

Captions for Movies S1 to S2

**Other Supplementary Materials for this manuscript include the following:**

Movies S1 to S2

#### Materials and Methods

##### Bacterial strains, media, and culture conditions

Relevant characteristics of bacterial strains, plasmids, and primers used in this study are listed in **Table S1, S2, and S3**. LB and M9 agar were used as complex and minimal media, respectively. Incubations were carried out at 37°C or 30°C for 16 hours (LB) to 48-72 hours (M9), except where noted. For plasmid maintenance and antibiotic selection markers, antibiotics (Sigma-Aldrich Pte. Ltd., Singapore) were added to the media at the following concentrations: ampicillin (100 µg/ml), kanamycin (50 µg/ml (for plasmid-based markers), 25 µg/ml (for chromosomal markers)), chloramphenicol (20 µg/ml), tetracycline (10 µg/ml), streptomycin (50 µg/ml), and gentamycin (3 µg/ml). For negative selection markers, media was supplemented with glucose or rhamnose, as indicated, both at 0.2% (Sigma-Aldrich Pte. Ltd., Singapore). Maintenance of plasmids carrying toxin genes was carried out in LB supplemented with 2% glucose. Minimal media was supplemented with nicotinic acid and lysine (Sigma-Aldrich Pte. Ltd., Singapore) for growth of *E. coli* UTI89 and TOP10, respectively.

All PCRs for cloning and recombination were carried out using Phusion high fidelity DNA polymerase. All diagnostic PCRs were carried out using DreamTaq DNA polymerase. All restriction enzymes were purchased from New England BioLabs, Singapore or Fermentas unless stated otherwise. All polymerases were purchased from ThermoFisher Scientific. Sanger sequencing was performed by AITbiotech, Singapore.

##### Molecular Cloning

**EZ-Tn5™ pMOD™-3 <R6Kγori/MCS> Transposon construction vectors**

**pSLC-390 (Ez-Tn5 pMOD-3 < $P_{\sigma 70}$-mCherry- $P_{rhaB}$-*tse2*-kan-oriT>)**

Markers were amplified by PCR and joined using an overlap-extension strategy. pSLC-130 was used as the template for amplification of $P_{\sigma70}$-mCherry, pSLC-389 as the template for $P_{rhaB}$-*tse2*, and rEc2 strain as the template for amplification of kan-*oriT* (**Table S3**). Primers were designed to have 30-35bp overlapping sequences to each fragment and an XmaI enzyme site at 5' ends. Agarose gel-purified overlapping PCR fragments were blunt ligated at the XmaI site into the pMOD-3 vector (cut with XmaI, dephosphorylated, and gel-purified). The ligation mixture was transformed into *E. coli* TOP10 competent cells and plated on LB agar with kanamycin. All incubations were carried out at 30°C, overnight and the clones were confirmed by phenotypic testing, PCR for the newly formed junctions between markers, and Sanger sequencing.

**pSLC-391** (*aadA* gene cloned with vsfGFP)

pSLC-294 was used as the template plasmid, which contains the vsfGFP-9 (*1*) gene cloned into pBAD33 vector. pSLC-294 was cut using the PsiI restriction enzyme, dephosphorylated, gel purified, and used as the vector. The *aadA* gene (conferring resistance to streptomycin and spectinomycin) was amplified from plasmid pAH144. The product was phosphorylated and blunt ligated into pSLC-294. The ligation mixture was transformed into *E. coli* TOP10 competent cells and plated on LB agar containing chloramphenicol, streptomycin, spectinomycin, and 0.2% glucose at 37 °C for 18-24 hours. Clones obtained were confirmed by phenotypic testing, PCR for the newly formed junctions, and Sanger sequencing.

**pSLC-392** (*chu* operon for complementation)

The *chu* operon was amplified from genomic DNA purified from wild type UTI89 and cloned into the pACYC177 vector using the BamHI and ApaLI restriction sites, disrupting the ampicillin resistance gene.

**Construction of donor and recipient strains**

Defined *E. coli* UTI89 Hfr donor strains were constructed by inserting the mCherry-$P_{rhaB}$-*tse2*-kan-*oriT* cassette from pSLC-390 amplified using primers with 50bp homology to the insertion site (**Table S3**), while MG1655 recipient strains were constructed using a similar strategy using the chlor-$P_{rhaB}$-*tse2* cassette from pSLC-281. The plasmids were cut with restriction enzymes ScaI (pSLC-390) and NotI (pSLC-281) for linearization before amplification and treated with DpnI, followed by agarose gel purification. All genomic manipulations were carried out using previously described Red recombinase recombineering protocols optimized for lab or clinical isolates of *E. coli* with minor modifications (*2*). pRK24, a conjugation helper plasmid, was introduced into the donor strains from Addgene strain no. 51949 using previously described conjugation procedures (*3*).

**Transposon mutagenesis**

For donor libraries, synthesis of a custom transposome was carried out by amplifying the mCherry-$P_{rhaB}$-*tse2*-kan-*oriT* cassette from pSLC-390 using primers ME plus 9-3' and ME plus 9-5' as suggested by the manufacturer (EZ-Tn5™ pMOD™-3 <R6Kγori/MCS> transposon construction vectors; **Table S3**). For the recipient library, synthesis of a custom transposome was carried out by amplifying the chlor-$P_{rhaB}$-*tse2* cassette from pSLC-281 using primers EZ-Tn5 pKD3 fwd and rev. These primers were synthesised with 5' phosphorylated ends and the PCR reactions were performed using Phusion High-Fidelity DNA polymerase (New England BioLabs). To assay for the stable regions of UTI89, a custom Tn5 transposon was created by PCR amplification of chlor-$P_{rhaB}$-*tse2* fragment from pSLC-281. Primers were designed to have 19bp mosaic repeats at their 5' ends. All amplified products were treated with the DpnI restriction enzyme. The gel-purified PCR product (1-3 μg) was mixed with 4 μl of Tn5 transposase to have a final reaction volume of 10 μl (the balance made up with 80% glycerol). This mixture was incubated at 37°C for an hour and stored at -20°C for up to one year.

Highly efficient competent cells of wild type UTI89 and MG1655 were prepared using standardized techniques. Briefly, Strains were grown overnight in 1-2 ml LB, diluted 1:100 the next day into fresh media and grown with agitation till $OD_{600}$ = 0.55–0.60. Cells were then chilled in an ice water bath for 30 min and harvested by centrifugation (5000 rpm for 10 min), washed three times with cold (4°C) water, re-suspended in 1/100 of the original culture volume of cold 10% glycerol, and frozen in 50 μl aliquots as competent cells. The whole procedure was carried out in the cold room. Transposon mutagenesis was carried out by electroporating 1 μl of a custom EZ-Tn5 transposome preparation (described above) into 100 μl thawed highly efficient competent cells using 1 mm electroporation cuvettes in a GenePulser XCEL (Bio-Rad, Singapore) set to an output voltage of 1700 V with 25 F capacitance and 400 Ω resistance. Cells were then recovered in pre-warmed LB with 2% glucose at 37°C with shaking for 2-3 h and plated on prewarmed LB agar supplemented with kanamycin or chloramphenicol (depending on the marker in the custom transposome) with overnight incubation at 37°C (except for transformations with *oriT*-containing transposomes, which were incubated at 30°C).

Following overnight incubation, the donor library for MAE was screened for mCherry positive clones using a fluorescent microscope (MVX10; Olympus, Singapore), followed by one round of purification (streaking to single colonies on LB-kanamycin supplemented with 2% glucose). The whole procedure was carried out three times to obtain a final donor library of 147 individual clones. The pRK24 helper plasmid was conjugated into each clone individually following previously published protocols (*3*), except that conjugations were performed in liquid cultures. Clones were then screened for stability of the transposon insertion (described below); stable clones were then pooled to make a final donor library for MAE. The recipient library for MAE was constructed by selecting 948 individual clones, which were also screened for stability of the transposon insertion (described below). 495 stable clones were then pooled together to make a recipient library. For MAE experiments, the recipient library was transformed with pSLC-391 to facilitate tracking of parents and transconjugants. For assaying the stable and unstable regions of the UTI89 chromosome, ~10,000 clones were pooled together as a sweep from the initial transformation without purification of individual colonies.

**Identifying clones with stable transposon insertions**

Individual donor and recipient colonies were grown in LB with 2% glucose, kanamycin, and tetracycline (for donor clones) or chloramphenicol (for recipient clones) overnight at 30°C or 37°C, respectively. The cultures were diluted in PBS with 10-fold serial dilutions and plated on non-restrictive (LB with 2% glucose and antibiotics, for total CFU) and restrictive (M9 with rhamnose) media; these plates were incubated at 37°C for 16 hours (non-restrictive agar) or 48-72 hours (restrictive agar). The loss rate of the custom transposon insertions was calculated as the ratio of the titer under restrictive conditions to that under non-restrictive conditions. Clones with a loss rate >=$10^{-5}$ were classified as "unstable", while those with loss rates < $10^{-5}$ were classified as "stable". Stable clones were pooled together to create the donor and recipient libraries for MAE.

To identify stable and unstable regions of the UTI89 genome, 20 µl of the UTI89 Tn5 library was thawed, diluted in PBS so as to get ~300 colonies on LB agar with chloramphenicol and 2% glucose. 1000 individual colonies were picked and resuspended in 50 µl PBS each, diluted with 10-fold serial dilutions in PBS, and plated on non-restrictive (LB with chloramphenicol and 2% glucose) and restrictive (M9 with rhamnose) solid media; these plates were incubated at 37°C for 16 hours (non-restrictive agar) to 48-72 hours (restrictive agar). The loss rate of the Tn5 insertions was calculated as the ratio of the titer under restrictive conditions to that under non-restrictive conditions. Clones with a loss rate >=$10^{-5}$ were classified as "unstable", while those with loss rates <$10^{-5}$ were classified as "stable". For sequencing, all 1000 individual clones were grown overnight in 500 µl LB broth with chloramphenicol and 2% glucose at 37$^{o}$C. 200 µl of each clone was pooled together into an "input" library (containing all 1000 clones). 200 µl of each clone was then also pooled together into a "stable" or "unstable" library, according to its classification. This experiment was carried out two times independently to screen a total of 2000 colonies.

**Conjugation**

*Directed transfers (generalized allelic exchange, GAE).* The conjugation protocol described by Natalie J Ma et al. (*3*) was modified to incorporate negative selection. Briefly, donor strains were grown overnight in LB with tetracycline, kanamycin, and 2% glucose at 30°C under shaking conditions. Recipient strains were grown in LB with chloramphenicol and 2% glucose at 37°C with shaking. Overnight cultures were diluted in 1:100 fresh LB with appropriate antibiotics and grown to $OD_{600}$ = 1.0 at appropriate temperatures with shaking. Cells were centrifuged at 5000 rpm for 8-10 mins, washed twice with 10 ml LB (no antibiotics) and resuspended to a final $OD_{600}$ = 20-25 in LB. The donor and recipient were mixed in a 4:1 ratio (total volume = 100 µl). This mating mixture was spotted on a pre-warmed LB agar plate and incubated at 30°C for 2.5-3 hrs. The mating mixture was then scraped off and resuspended in 750 µl of PBS. Another 750 µl of PBS was added to this culture, which was then centrifuged at 13,500g for 1 min. The pellets were resuspended in 200 µl PBS. Ten-fold serial dilutions were plated on LB with antibiotics (control plates for total counts) and M9 plates with rhamnose (without niacin) for negative selection. All plates were incubated at 37°C for 16-20 hrs (LB plates) or 60-72 hrs (M9 plates). As controls, donor cells (80 µl) and recipient cells (20 µl) were also treated exactly the same way as conjugation mixtures and plated on control plates as well as test plates. Transconjugants were confirmed by phenotypic and genetic testing (PCR and Sanger sequencing).

*Mass allelic exchange*. The protocol was similar to the GAE protocol above with minor modifications and larger volumes. 50 µl of the donor library was thawed and grown overnight in 5 ml LB with tetracycline, kanamycin, and 2% glucose at 30°C under shaking conditions. 50 ul of the recipient library was grown in 5 ml LB with streptomycin, spectinomycin (for plasmid pSLC-391), chloramphenicol, and 2% glucose at 37°C with shaking. The rest of the procedure remained the same as that described above for GAE. Transconjugants were selected using a combination of positive and negative selection on M9 rhamnose plates without niacin and supplemented with streptomycin and spectinomycin. To maximize the number of transconjugants, conjugation mixtures were plated on 180 test plates and 1 control plate each. All plates were incubated at 37°C for 16-20 hrs (control LB plates) or 60-72 hrs (M9 plates).

Following incubation, all GFP positive colonies in dilutions over the recipient background were pooled together and stored as transconjugant libraries. Donor and recipient controls were maintained as described above. This whole procedure was carried out two times to get two independent transconjugant libraries.

**Serum resistance assay**

The pooled transconjugant library was thawed and diluted to a concentration of ~$10^3$ CFU / 50 µl. 50 µl were seeded into each well of a 96-well plate and 50 µl of undiluted human serum (Sigma-Aldrich Pte. Ltd., Singapore) was added to each well. The plate was then incubated at 37°C with shaking at 15 rpm for 3 h. After serum treatment, the serum + culture was removed and fresh LB (150 µl) was added to the wells with overnight incubation at 37°C. Single clones were streaked out from wells that showed growth after serum treatment; these individual clones were then retested twice to verify serum resistance. Wild type MG1655 and UTI89 were included on each plate as negative and positive controls, respectively.

For testing of single clones, each colony was grown overnight in LB at 37°C, then ~$10^3$ CFU were seeded into a single well. 50 µl of undiluted human serum was added and the reactions were incubated at 37°C at 15 rpm for 3 h. Serial dilutions of a 50 µl aliquot at 0 h (input) and at 3 h (output) were titered on LB agar.

**Phage cross-streak assay**

A cross-streak assay was used to test for resistance to P1*vir* and K1 bacteriophage. 50 µl of the phage lysate was streaked through the middle of an LB plate containing 2.5 mM $CaCl_2$ while overnight test cultures diluted to $OD_{600}$ = 0.5 were streaked perpendicular to the lysate. The plates were incubated at 37°C for 12 hrs and examined for phage resistant clones. Clones were classified as sensitive to the phage if there was a zone of inhibition around the phage lysate; uniform growth through the streak was classified as resistance. Wild type MG1655 and UTI89 were used as positive (sensitive) and negative (resistant) controls, respectively.

**Phage lysis assay**

Overnight cultures were diluted 1:10 in 6 ml LB and incubated at 37°C under shaking conditions. When the $OD_{600}$ reached 0.25-0.30, 50 µl of phage lysate ($10^9$ or $10^{10}$ PFU/ml) and 50 µl of 1M $CaCl_2$ was added to the culture, and incubated at 37°C for 3-4 h with shaking. Phage resistance and sensitivity were assessed by the lysis/turbidity of the culture, as well as by titering the culture on LB agar before and after phage treatment.

**Whole Genome Sequencing**

Genomic DNA for whole genome sequencing was extracted using either the Wizard Genomic® DNA Extraction Kit (Promega), QIAmp DNA mini kit (Qiagen, Singapore) or Promega Wizard® SV 96 genomic DNA purification system according to manufacturers' protocols. DNA was eluted in 50-100 µl of nuclease free water and quantified using Qubit 2.0 fluorometer (Invitrogen) and Qubit dsDNA high sensitivity kit (Life Technologies; Q32854).

Whole genome sequencing libraries were prepared using either Illumina TruSeq DNA Nano library preparation kit (where maximum of 24 samples were multiplexed together) or Illumina Nextera DNA library prep kit (where 96 to 384 samples were multiplexed together) according to the manufacturer's recommendations. Sequencing was done on an Illumina HiSeq4000 or NextSeq with 2 × 151 bp reads depending on machine availability.

Raw fastq files were mapped using bwa (version 0.7.17) to the MG1655 (NC_000913.3) genome. SNPs were called using lofreq* (version 2.1.2). SNPs were filtered for those that were found in at least 7/8 independently sequenced libraries of UTI89 similarly mapped to the MG1655 reference sequence. Recombination blocks were identified as the minimal region spanning consecutive SNPs found in the control UTI89 libraries. We set a minimum cutoff of 16 Kb for recombination blocks to analyze the statistics of transferred regions.

**Transposon-Directed Insertion Site Sequencing (TraDIS)**

TraDIS was performed similarly to previous reports with minor modifications (*4*). Briefly, gDNA was diluted to a concentration of 100 ng in 52.5 µl of nuclease free water and sheared using a Covaris® S2 machine as per the manufacturer's recommendations for the Illumina TruSeq nano DNA library prep protocol. DNA end repair, DNA 3' end adenylation, and adaptor ligation were carried out using the Illumina TruSeq low sample protocol. After adapter ligation, enrichment of adapter-ligated fragments was performed using a custom transposon-specific indexing forward primer and the Illumina reverse primer Index I (**Table S3**). The forward primer was designed based on (*4*) which is a 89 bp primer with 25 bp specific to the custom Tn5-transposons (25 bp) and carrying a 64 bp overhang which includes a 6 bp index sequence (from Illumina adaptors sequences), the 33 bp binding site for the Illumina Read 1 sequencing primer, and the 23 bp P5 sequence for binding to an Illumina flow cell. Amplification was done using the KAPA library amplification kit (KAPA biosystems) using the following reaction mixture: 25 µl

2× KAPA HiFi hot start ready mix, 2.5 µl 500 nM forward primer, 2.5 µl 500 nM reverse primer, and 20 µl library DNA. The enrichment PCR thermocycling protocol was: 98$^{o}$C for 45 sec; 22 cycles of 98$^{o}$C for 15 sec, 60$^{o}$C for 30 sec, and 72$^{o}$C for 30 sec; and a final extension at 72$^{o}$C for 1 min. PCR cleanup was done as per the Illumina TruSeq Nano protocol. Agilent DNA 1000 chips were used on Agilent Technology 2100 Bioanalyser for quantification and quality assessment for all libraries. Libraries were normalized to 10 nM in a total volume of 30 µl and 5 µl of each library was pooled together for sequencing on an Illumina NextSeq 500 system with 2 × 151 bp reads sequencing.

Fastq reads were demultiplexed a second time using deML (version 1.1.3), with the expected transposon sequence and internal barcode used as custom demultiplexing sequences for the R1 reads. The remaining R1 read sequence was used with the corresponding R2 reads for mapping to the MG1655 (NC_000913.3) or UTI89 (NC_007946.1 and NC_007941.1) genome sequence using bwa mem (version 0.7.17) with default parameters. The corresponding bam file was filtered to include only properly mapped pairs (as reported by bwa). The insertion location was then extracted; a canonical "left insertion position" was calculated by correcting the position for R1 reads mapped to the negative strand by subtracting 9 bp from the reported mapping coordinate of the 5' end of the original read.

**Saponin-facilitated pod formation**

Bladder epithelial cell line 5637 (ATCC HTB-9™) was used to carry out *in vitro* pod formation as previously reported (*5*). RPMI 1640 medium (Gibco, Singapore) supplemented with 10% fetal bovine serum (Gibco, Singapore) was used for cultivation and all incubations were carried out at

37°C with 5% $CO_2$. All bacterial strains used in these experiments carried pSLC-391 (expressing vsfGFP-9) for pod visualization and were cultured under Type 1 pilus-inducing conditions (2 × 24 h passages in LB under static conditions at 37°C, with a 1:1000 dilution between passages) (*6*). 5637 cells were seeded in 6-well, 12-well, or 96-well (tissue culture treated CellCarrier-96 ultra micro plates) plates with 70% confluency to get 100% confluency in 24 h. Within 24 h, the cells were infected with controls (UTI89 and MG1655), individual transconjugants, or the pooled transconjugant library at an MOI (multiplicity of infection) of 10 bacteria per host cell. Following inoculation, plates were centrifuged at 600× g for 5 min to synchronize bacterial contact with the epithelial cells. These plates were then incubated at 37°C with 5% $CO_2$ for 2 h for infection to occur. Then, gentamycin was added to a final concentration of 100 μg/ml to kill any extracellular bacteria and incubated again at 37°C with 5% $CO_2$ for 2 h. Cells were then washed with PBS supplemented with $Mg^{2+}$ and $Ca^{2+}$ and overlaid with RPMI (10% FBS) with gentamycin at 10 μg/ml for incubation at 37°C with 5% $CO_2$ overnight. Host cells were washed 3 times with PBS supplemented with $Mg^{2+}$ and $Ca^{2+}$ after overnight incubation and incubated again at 37°C with 5% $CO_2$ for 15 mins with fresh media containing 0.025% saponin (no saponin as control). After 30 min, cells were washed again with PBS supplemented with $Mg^{2+}$ and $Ca^{2+}$ and incubated for another 4h (UTI89) to 6h (MG1655 and transconjugants) to allow bacterial growth. The resulting cells were examined microscopically to look for intracellular pod formation.

**Isolation of single pod-containing epithelial cells**

After 6 hr of saponin treatment, individual pod-containing 5637 cells were visualised under a fluorescence microscope (MVX10; Olympus, Singapore) and isolated by mouth pipetting, using a protocol previously described for isolation of single intracellular bacterial communities in a mouse model of UTI (*7*).

**Imaging pods for quantification**

96-well (tissue culture treated CellCarrier-96 ultra micro plates) plates were used for visualization and quantification of pods using an Opera Phenix High-Content Screening System or an Operetta CLS High-Content Analysis System (PerkinElmer). After 4-6 hours of saponin treatment, the cells were fixed with paraformaldehyde (final concentration to 2%) at 37°C under 5% $CO_2$ for 20-30 mins. After fixation, cells were stained with DAPI (1:1000 in $DPBS^{2+}$) and WGA (5 μg/ml in $DPBS^{2+}$) at 37°C for 10-15 mins and visualised using 3 channels (Alexa 594 (WGA 594), DAPI, and EGFP) under a 40x water objective lens with confocal settings.

For live imaging, cells were incubated at 37°C with 5% $CO_2$ for 1 hour, after which the plates were transferred to a PerkinElmer Operetta CLS machine maintained under the same conditions (37°C, 5% $CO_2$). Each well was imaged using the non-confocal 40x air objective using brightfield and EGFP filters at 1 hour time intervals over 8-9 hours. Images were analysed using the Harmony high-content image and analysis software (PerkinElmer) and combined using ImageJ to create time lapse videos.

#### Supplementary Text

##### Supplementary Text 1 - Loss of negative selection stringency using transposon libraries

We made recipient and donor libraries by mutagenesis using a custom transposon carrying the negative selection cassette. Initial tests of the negative selection stringency for these libraries showed a very high frequency of growth when plated on rhamnose ($10^{-3}$). This frequency was several orders of magnitude higher than those previously reported for individual insertion locations ($10^{-7}$). We hypothesized that, because some regions of the genome are inherently unstable (such as plasmids, pathogenicity islands, prophages, and other mobile genetic elements) (*8*, *9*), clones with the negative selection cassette inserted in these were contributing to the low negative selection stringency of the libraries. Indeed, when we inserted the negative selection cassette into the PAI-II pathogenicity island in UTI89, we measured a similarly high frequency of rhamnose resistant colonies ($10^{-3}$; **Fig. S1A**). To surmount this problem, we screened the donor and the recipient transposon libraries for clones with stable insertions of the transposon (defined as < $10^{-6}$ CFU/ml when titered on rhamnose); these "stable clones" were then pooled to make the final libraries used for conjugation.

As a side product of this screening, we also had a pool of "unstable clones", which allowed us to directly identify unstable regions of the *E. coli* genome. This application was again feasible due to the advance in the selection stringency added by our negative selection system. We carried out another transposon mutagenesis in wild type UTI89, using the same custom transposon ($P_{rhaB}$-chlor-*tse2*) as the one used to engineer the recipient libraries used for MAE. Using negative selection, two thousand individual colonies from this library were screened to differentiate between clones that had stable vs unstable Tn insertions. Approximately 20% of the insertions were unstable. We performed TraDIS (transposon-directed insertion site sequencing) by pooling all the stable and unstable insertions in two distinct libraries and as expected, we found prophages, PAIs, and the plasmid, but also some novel unstable regions that may provide new insights into mechanisms of chromosome plasticity, maintenance, and evolution (**Fig. S1B**).

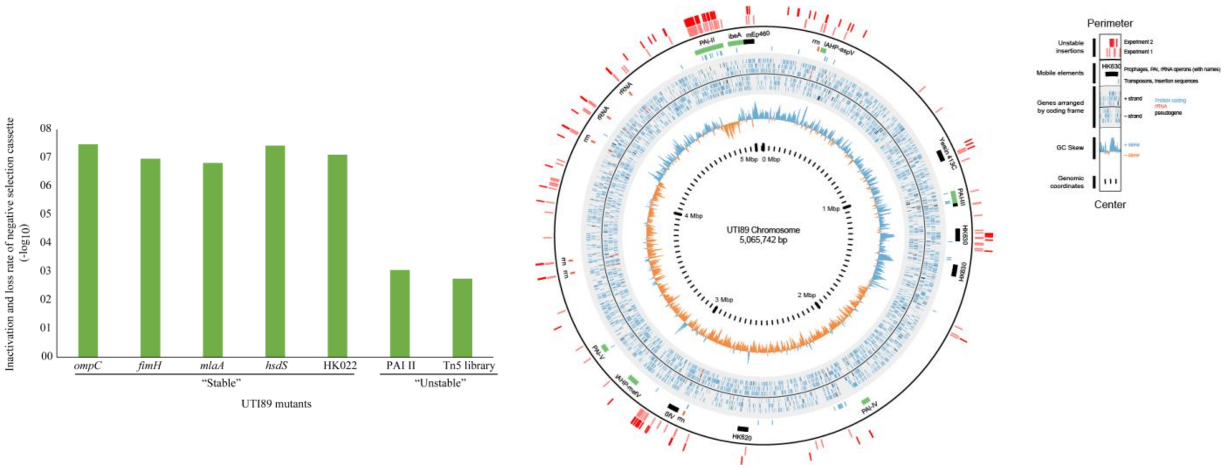


**Fig. S1A.** Characterization of genome instability in UTI89. (**A**) Inactivation and loss rate of the negative selection cassette in UTI89. Insertion site of the negative selection cassette determines its loss rate/inactivation. Bars represent negative selection stringency (CFU/ml on restrictive agar over CFU/ml on non-restrictive agar) of the dual selection cassette at different insertion sites on the chromosome and of a UTI89 transposon library. Stable and unstable genes/locations are marked. PAI II refers to Pathogenicity Island II. (**B**) Map of chromosomal instability in UTI89. The data plotted in each track is indicated in the legend at the top right. Rings, from inside to outside, are GC skew, annotated genes with gray background, mobile elements, then insertion locations for unstable clones in two experiments. Genes are indicated as negative strand on the inner three circles, positive strand on the outer three circles, with colors indicating type of gene: blue = protein coding, red = rRNA, black = pseudogene. For mobile elements, phages (as called by Phaster) are black, rRNAs are red, and transposon sequences / insertion sequences are blue.

#### Supplementary Text 2 - Bias in the donor, recipient, and transconjugants libraries

We noted a clear bias for transferred regions near the origin of replication in the selected hybrids (**Fig S2A**). To investigate the source of this bias, we pooled clones with stable Tn insertions from the donor and the recipient libraries and carried out TraDIS to map their insertion sites. The donor pool consisted of 106 clones, while the recipient pool consisted of 495 clones. We found that unique insertion locations were biased towards the replication origin (*oriC*) in both pools (**Fig. S2B, S2C**). This correlates with the read depth per insertion and is consistent with other reports of transposon insertion bias due to the higher copy number of origin-proximal DNA (*10*). We observed an approximately 3x and 7x higher maximum insertion frequency in the donor and recipient libraries, respectively. We would expect an additional round of amplification of this bias during the conjugation process itself; we thus calculate an expected bias in the transconjugants of 63-147x. This is not significantly different from our observation of only 1 clone with a conjugation outside the origin-proximal region in 384 total sequenced clones; this suggests that the MAE process itself, using conjugation and selection against the negative selection markers, does not inherently introduce additional bias.

To verify this last suggestion, we directly tested the efficiency of hybridization using directed transfers. We selected individual donor and recipient clones with insertions of the *oriT* and negative selection transposons, respectively, near the origin and the terminus (**Fig. S2D**). In these control hybridizations, we saw no trend towards lower efficiency in the terminus-proximal transfers (**Fig. S2E**). This data confirm that there is no general bias against hybridization in the terminus-proximal region of the recipient chromosome.

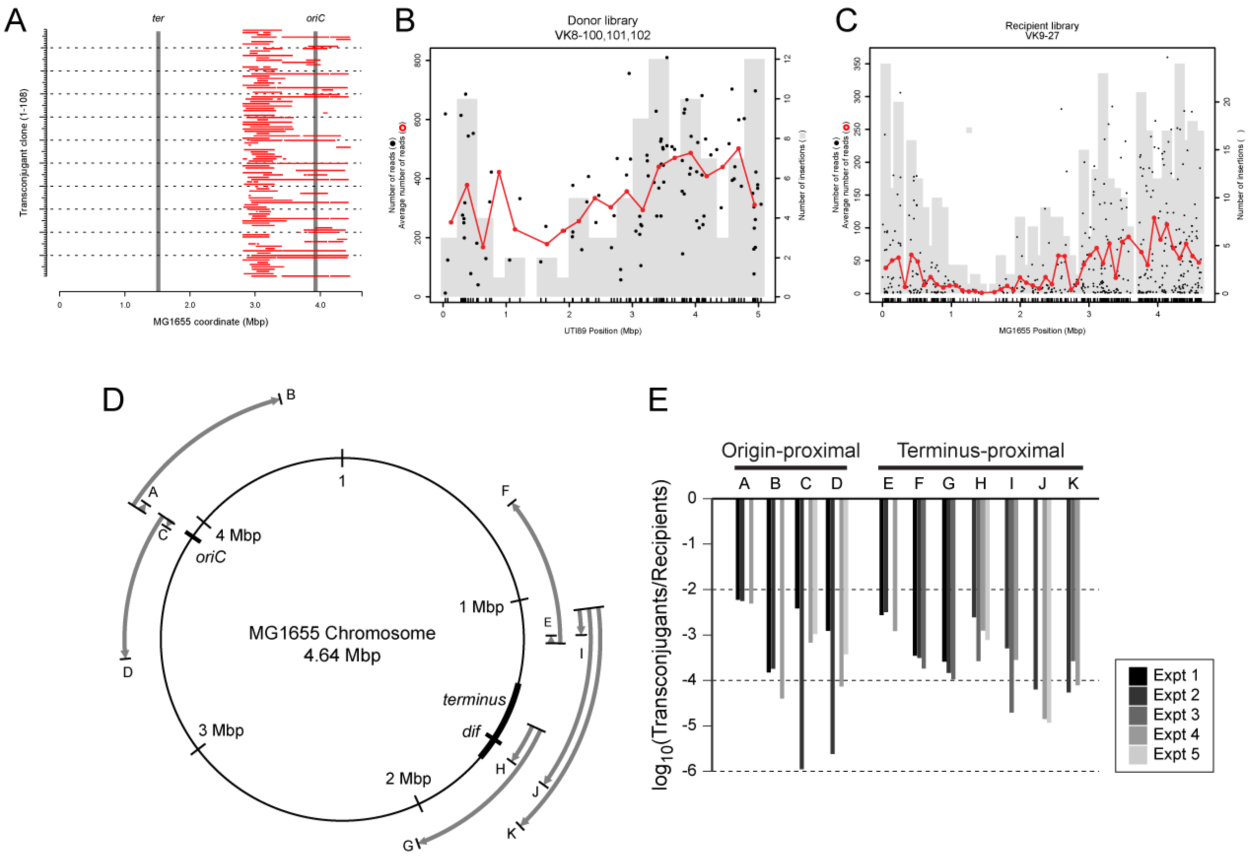


**Fig. S2. (A)** Transferred SNPs for 108 individual clones from one hybridization. Each clone was analyzed by whole genome sequencing, using the MG1655 genome as a reference. Genomic coordinate is plotted on the x-axis. Individual clones are plotted at different locations on the y-axis. Blocks of consecutive single nucleotide polymorphisms (SNPs) relative to the MG1655 genome sequence (corresponding to UTI89-specific sequence) are represented by red bars. Thick vertical gray lines indicate the positions of the origin (*oriC*) and terminus (*ter*). (**B**) Insertions positions for the customized *oriT*-containing transposon in the donor library. (**C**) Insertion positions for the customized negative-selection cassette-containing transposon in the recipient library. Insertion locations were obtained from a TraDIS experiment on the recipient library. In all, 461 unique insertion locations were identified from a pooled library of 495 clones. The individual insertion locations are indicated by the short vertical black lines directly above the x-axis. Black dots indicate the number of reads supporting each insertion site in the TraDIS sequencing, using the left y-axis. The gray vertical bars represent a histogram of the number of unique insertion sites, using the right y-axis. The red open circles plot the average read depth for all insertions within each histogram bin, using the left y-axis. (**D**) Schematic of directed transfers. The MG1655 chromosome is represented by the central complete circle. The genomic coordinates are indicated by the numbered tick marks. The replication origin (*oriC*) and terminus region (including the *dif* site) are indicated by thicker black lines. 11 directed transfers are indicated, lettered A-K. For each, a black line at the beginning of a gray curved arrow indicates the position of the *oriT* inserted into the donor (UTI89), represented at the homologous location in the recipient (MG1655). A black line at the end of a gray curved arrow indicates the location of the negative selection cassette in the recipient, which should be minimally spanned by a successful recombination. The thicker gray line and the arrowhead indicate the direction of transfer of DNA from the donor to the recipient. (**E**) Efficiencies of transfer. The efficiencies of transconjugants resulting from the directed transfers indicated in (C) are shown as a bar graph. Each bar represents the number of resulting transconjugants divided by the number of recipients that were put into the conjugation mixture, graphed on a logarithmic scale; transconjugants were verified by testing for the predicted lack of resistance to chloramphenicol and kanamycin, and in limited cases by Sanger sequencing of a locus near the insertion location of the negative selection cassette in the recipient. Seven of the transfers were repeated a second time (biological replicate); these are shown as separate bars.

#### Supplementary Text 3 - Statistics and distribution of transferred segments

We performed whole genome sequencing on 386 randomly selected transconjugant clones across 3 separate hybridizations. After quality control for sequencing depth and quality, we had 353 total transconjugant strains. Based on SNPs between UTI89 and MG1655, we then mapped the length of the transferred (recombined) blocks. All lengths are measured in terms of the original MG1655 coordinates from the first to the last SNP. To understand the statistics of large chromosomal transfers, we excluded recombination blocks below 16 Kb for the following analysis.

The majority (267, 73.6%) of the transconjugants had only one detectable recombination block. One strain each had 4 and 5 recombination blocks (**Fig. S3A**). The distribution of the length of the recombined regions was trimodal, with several large (> 1 Mbp) transfers, a second peak at ~400 Kb, and a third peak at ~50 Kb (**Fig. S3B**). Strains with multiple recombined regions must have had at least one recombination whose location was not determined by the transposon containing the negative selection marker in the recipient. Given that the smaller recombination blocks seem to be occurring in strains that had more than one recombined region, we asked whether recombination blocks in such strains were systematically biased towards shorter lengths. We compared the length distribution of the larger recombination block in strains with 2 transfers with the length distribution of the sole recombination block in strains with one transfer (**Fig. S3C**). The medians of these distributions were not statistically different ($P=0.9242$, two-sided Wilcoxon rank sum test). This suggests that in strains where more than one recombination has occurred, at least one of the recombinations (perhaps a "primary" recombination) has the same recombination statistics as strains where only one recombination has occurred, and we suspect that such primary recombination events are directed by the negative selection cassette.

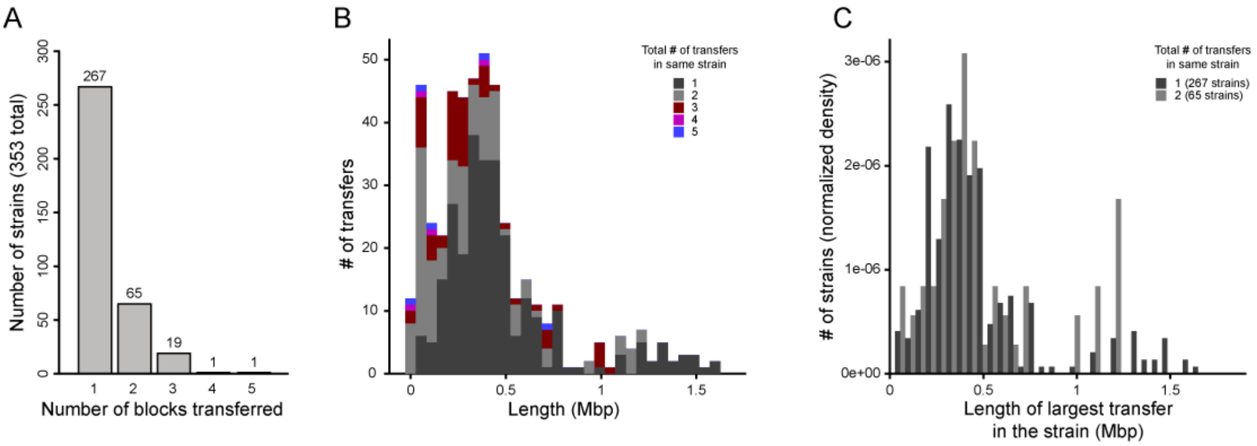


**Fig. S3.** Statistics of recombination blocks in transconjugants. (**A**) Histogram of the number of recombination blocks per strain. The number of strains (y-axis) is plotted on the y-axis; the number of recombination blocks is plotted on the x-axis. Small numbers above the bars indicate the height of the bar. (**B**) Histogram of the length of recombination blocks. All recombination blocks greater than 16 Kb were included in this analysis. The number of recombination blocks is plotted on the y-axis; the length of the recombination blocks is indicated on the x-axis. The color of the bars represents the total number of recombination blocks in the strain carrying a recombination block (thus, all recombination blocks colored blue (5 total recombination blocks) were present in the same transconjugant clone). (**C**) Comparison of the length distribution of the longest recombination block in strains with 1 (black) or 2 (gray) recombinations. The normalized density (arbitrary units) is plotted on the y-axis; the total height of the bars for each color is the same. The length of the recombination block is plotted on the x-axis.

**Supplementary Text 4 - Confirmation of serum and phage resistance phenotypes in isolated transconjugants**

The pooled transconjugant library containing ~12,000 hybrids (VK10-85) was screened for clones that had gained the serum resistance phenotype using human serum. We isolated 60 clones and repeated the serum treatment individually on these clones to confirm the phenotype (**Fig. S4A**). MG1655 and a randomly selected serum sensitive clone (green “S”) were used as negative controls, and UTI89 was used as a positive control. Since the serum and P1 phage sensitivity is known to be a single locus phenotype in MG1655, we also confirmed the P1 resistance of these clones using phage streak assays (**Fig. S4B**). As expected, all 60 clones that were serum resistant were also phage resistant, while the serum sensitive clone was sensitive to P1 phage (data not shown).

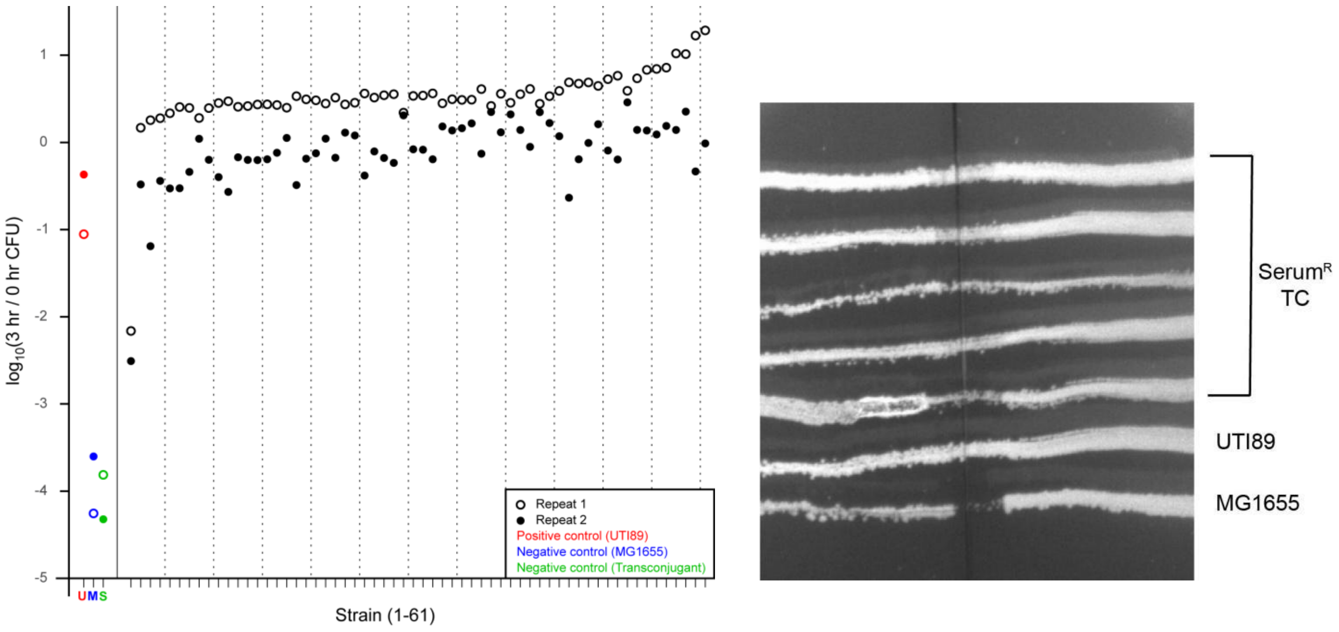


**Fig. S4A.** Validation of serum sensitivity of transconjugants. 60 resistant clones and one sensitive clone (as a control) isolated from the pooled serum sensitivity screen were retested individually. CFU titers for each clone were measured before (0 hours) and after (3 hours) treatment with serum. The logarithm of the ratio of these CFU titers is plotted on the y-axis; different clones are plotted on the x-axis. Open and closed symbols represent two different biological repeats of the same experiment. UTI89 was used as a positive control and is indicated by red symbols and a red “U” on the x-axis. MG1655 was used as a negative control and is indicated by blue symbols and a blue “M” on the x-axis. The sensitive clone is indicated by a green “S” on the x-axis. A solid vertical line separates the controls from the resistant transconjugants; dotted vertical lines are placed every 5 clones to help with visualization. (**B)** Phage cross streak assay for confirmation of P1 phage resistance for a few representative strains. P1 phage was streaked across the center of the plate and individual bacterial cultures to be tested were streaked perpendicular to the phage streak. Zone of inhibition around the streak (MG1655)

represents phage sensitive clones while uniform growth through the phage (UTI89) represents resistant clones. TC = transconjugants

**Supplementary Text 5 - Transfer of the *rfb* locus from UTI89 to MG1655 confers serum and P1 phage resistance**

To confirm the serum and phage resistance phenotype observed using MAE, we verified that a targeted transfer of the entire *rfb* locus from UTI89 to MG1655 indeed confers these phenotypes. A UTI89 donor strain (with *oriT* inserted upstream and directed towards the *rfb* cluster) was crossed with MG1655 (with the selection cassette downstream of the *rfb* cluster). All other experimental conditions were identical to the MAE experiments. 34 single transconjugants were isolated and screened for resistance to human serum and P1 phage. 13 transconjugants were confirmed to be resistant to both serum and P1 phage (**Fig. S5A and S5B**). Illumina whole genome sequencing was carried out on 4 serum$^{R}$ phage$^{R}$ and 6 serum$^{S}$ phage$^{S}$ clones of this experiment. As expected, we observed staggered fragments of the donor chromosome (of varying lengths) recombined into the recipient chromosome, leading to the inclusion or exclusion of the *rfb* cluster in different transconjugants. All serum$^{R}$ phage$^{R}$ transconjugants had transferred *rfb*$^{UTI89}$; in contrast, all serum$^{S}$ phage$^{S}$ transconjugants retained *rfb*$^{MG1655}$ (in the latter transconjugants, the incoming donor DNA recombined upstream of the cluster).

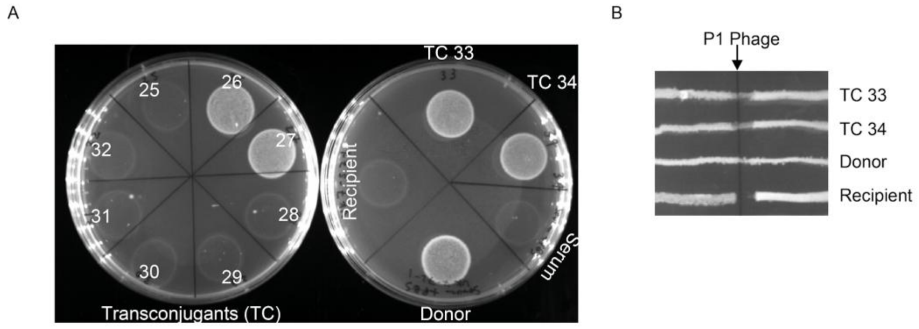

**Fig. S5**. Directed transfer of the rfb locus from UTI89 into MG1655 confers serum and P1 phage resistance. (**A**) Representative strains confirming resistance to serum. Single transconjugants were treated with serum and plated on LB to look for survival. Growth on LB represents serum resistance. (**B**) Representative strains confirming resistance to P1 phage. P1 phage was streaked across the center of the plate and individual bacterial cultures to be tested were streaked perpendicular to the phage streak. Zone of inhibition around the streak represents phage sensitive clones while uniform growth through the phage represents resistant clones. For both assays, the recipient strain (MG1655) was used as the negative control, while the donor strain (Hfr UTI89) was used as the positive control. Serum (without bacteria) only control was also plated. TC = transconjugant.

**Supplementary Text 6 - The *chu* operon is sufficient to confer formation of large pods to MG1655**

We used saponin-mediated pod formation assays on individual strains to validate the results of the screen. For all strains, technical replicates (i.e. replicate plates) were analyzed by time lapse video microscopy of unfixed cells (hourly for 8-12 hours) and by fluorescence microscopy of fixed cells. Pod area in pixels was measured from images of fixed cells. As previously reported, wt UTI89 formed large pods (> 10,000 pixels at 400x magnification) (**Fig. 3D, Fig. S6B**); a video of a representative large pod is shown in **Movie S1**. Wild-type MG1655 formed no such large pods (**Movie S1, Fig. 3D, Fig. S6B**). We noted that both strains formed smaller collections of bacteria, some of which appeared intracellular but did not expand like typical pods formed by UTI89; such structures were easily distinguished by microscopy of fixed cells or by time lapse video (**Movie S1; red blocks, Fig. S6B**).

Using these assays, we then verified that introduction of the *chu* operon into wt MG1655 (but not the empty vector) conferred the ability to form large pods to MG1655 (total pod numbers shown in main **Figure 3E**; a representative video is shown in **Movie S1, right**). Importantly, in VK9-57-11, a knockout of the *chu* operon abolished pod formation, a phenotype that was restored by complementation (**Movie S2, Fig. S6A and S6C**). Consistent with previous reports, a knockout of the *chu* operon in UTI89 decreased the size of the pods formed; in addition, by video analysis, these appeared to be smaller due to delayed kinetics of pod formation (**Movie S2 and Fig. 6C**).

Notably, we found that the saponin-mediated pod formation assay was highly variable between biological replicates (**Fig. S6D**). This was not due to cell passage number, as we tested multiple (independently purchased) stocks of 5637 cells from ATCC. We attempted to perform automated image analysis to identify pods, but the variability in the UTI89 and MG1655 controls led to the training algorithms identifying small bacterial collections that were qualitatively unlike the large pods described in (Mulvey paper) and observed in some replicates of our UTI89 controls (and none of the MG1655 controls) (**Fig. S6A**). Therefore, we set a minimum area cutoff of 10,000 pixels to define "large" pods, and all assays were performed with technical duplicates; one was analyzed by automated image processing of fixed cells, while the other was analyzed qualitatively by video microscopy of unfixed cells. The results of both the quantitative automated identification of large pods (**Figure 3E**) was in agreement with the qualitative video microscopy analysis (**Movies S1 and S2**).

**Videos 1 and 2.** Each frame shows a montage of all fields that cover an entire well of a 96-well plate. Strains are indicated at the bottom. The time in hours is indicated at the top left. Scale bar is indicated at the bottom left. All pixel values were normalized uniformly across all fields, wells, and time points to the same maximum pixel value, with a gamma of 2.0. The maximum pixel value was set at 90th percentile value for the brightest well (across all time points for that well) in each video.

**Fig. S6.** (**A**) Representative images of the pod-forming assay. Each strain is represented by three images; the images on the left are the DAPI (blue) and wheat germ agglutinin (orange) channels, middle are GFP (green) channel. The right image is a merger of all channels. (**B**) Expanded view of individual fields from Video 1. Left, green (GFP) channel; right, brightfield. Strains are indicated by the labels at the top, time in hours on the left. The pixel values are normalized to the full range for each image for each strain separately. The scale bar is the same for all images and indicated at the bottom left. (**C**) Expanded view of individual fields from Video 2. Left, green (GFP) channel; right, brightfield. Strains are indicated by the labels at the top, time in hours on the left. The pixel values are normalized to the full range for each image for each strain separately. The scale bar is the same for all images and indicated at the bottom left. (**D**) As shown, over 8 replicates, there was a bimodal behavior of the control strains, which was also seen in the test transconjugants. The pod identification was tuned on a preliminary experiment which gave similar results to the first three experiments, and the parameters were not changed during the analysis of any of the 8 experiments. Overall cell density and cells analyzed was not different (graph of cells, area, nuclei). There could have been some differences in the host cells, as the passage number was changing with every experiment. We therefore analyzed experiments 1-3 and 4-8 separately. Results for aggregated results from experiments 1-3 are shown here.

(A)

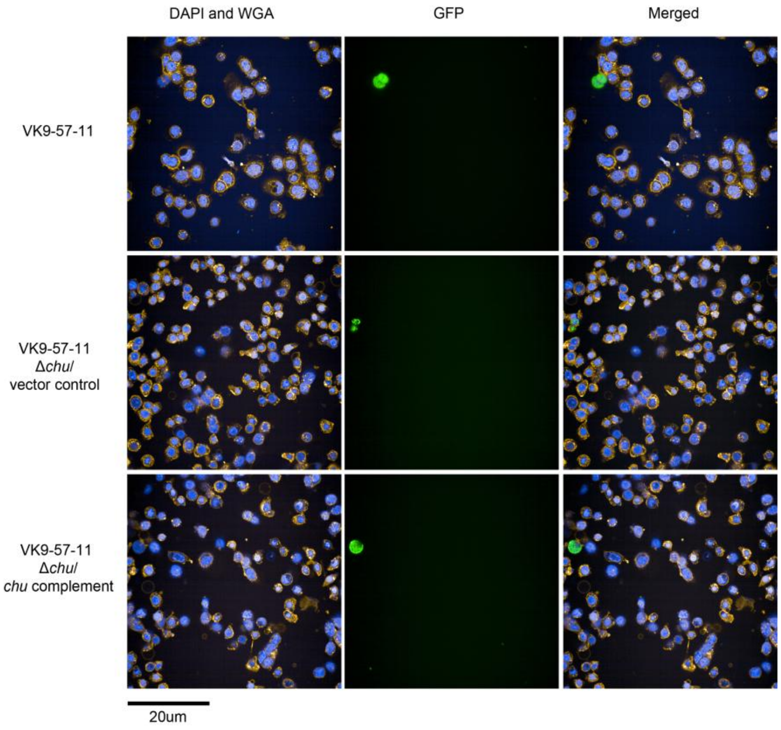

(B)

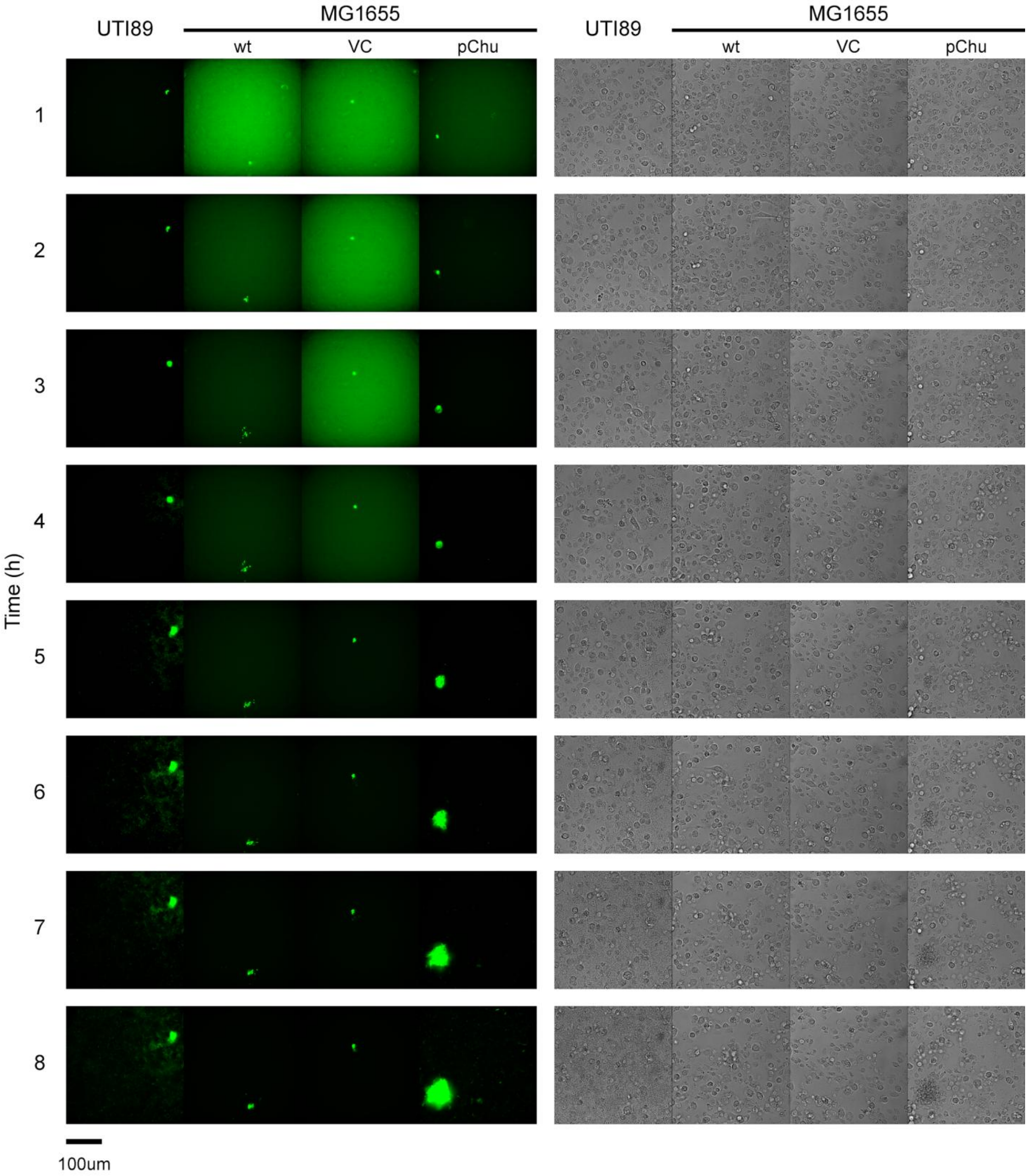
UTI89
MG1655
wt
VC
pChu
UTI89
MG1655
wt
VC
pChu
1
2
3
4
5
6
7
8
Time (h)
100um

(C)

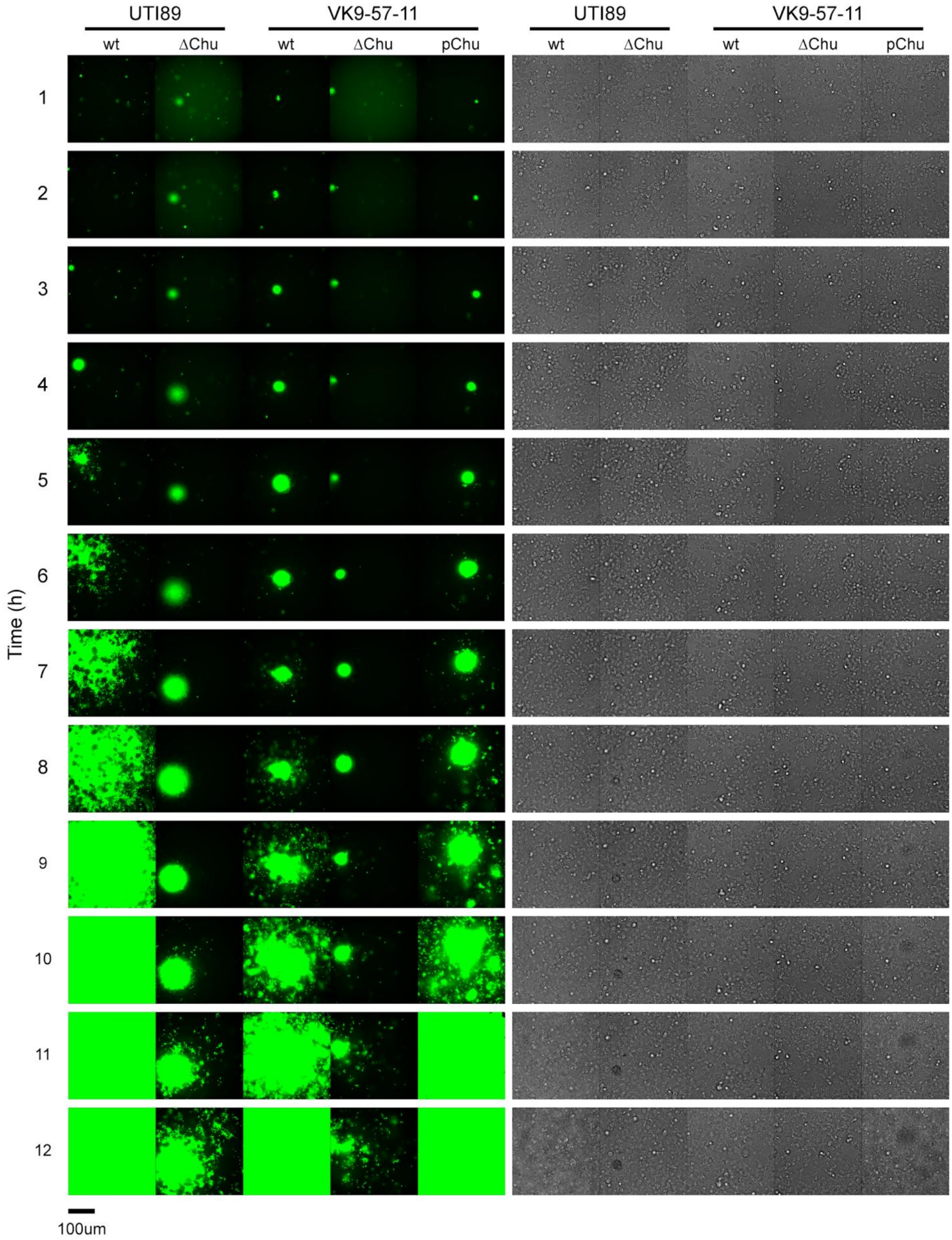
UTI89
VK9-57-11
UTI89
VK9-57-11
wt
ΔChu
wt
ΔChu
pChu
wt
ΔChu
wt
ΔChu
pChu
Time (h)
1
2
3
4
5
6
7
8
9
10
11
12
100um

(D)

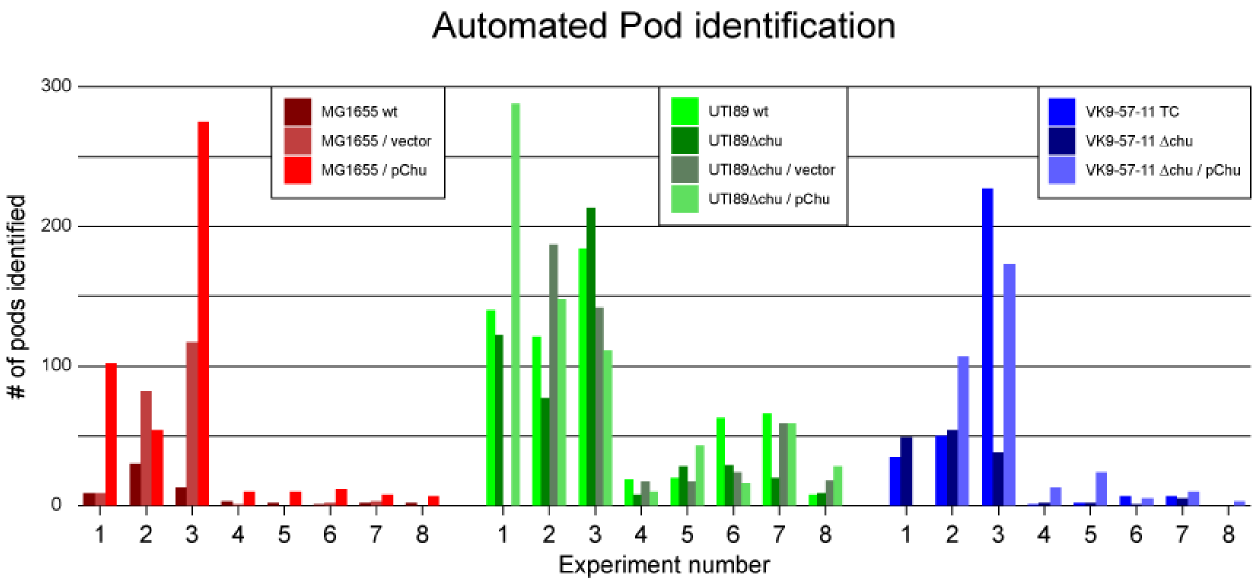

Automated Pod identification
### of pods identified
300
200
100
0
MG1655 wt
MG1655 / vector
MG1655 / pChu
UTI89 wt
UTI89Δchu
UTI89Δchu / vector
UTI89Δchu / pChu
VK9-57-11 TC
VK9-57-11 Δchu
VK9-57-11 Δchu / pChu
1 2 3 4 5 6 7 8
1 2 3 4 5 6 7 8
1 2 3 4 5 6 7 8
Experiment number

**Table S1. Bacterial strains**

| Strains | Genotype and/or relevant characteristics | Source/ reference |
|---|---|---|
| BW23473 | | Haldimann et al.,1997 |
| *E. coli* TOP10 | | |
| *E. coli* MG1655 K-12 | | |
| *E. coli* UTI89 | Clinical isolate | Chen et al., 2009 |
| rEc2 | Bacterial strain: 51946<br>kan-*oriT* template | Addgene/ Natalie J Ma et al., 2014 |
| SLC-557 | UTI89 *ompC*::kan-P*rhaB*-*relE* | Khetrapal et al., 2015 |
| SLC-1221 | UTI89 PAI II::kan-$P_{rhaB}$-*relE* | This study |
| VK5-102 | UTI89 Tn5 library ME-chlor-$P_{rhaB}$-*tse2*-ME | This study |
| UTI89 Donor library VK6-74-1 to 32<br>VK6-89-1 to 63<br>VK7-69-1 to 70 | UTI89 Tn5 library ME-$P_{\sigma 70}$-mCherry- $P_{rhaB}$-*tse2*-kan-oriT-ME | This study |
| Pooled donor library for MAE VK8-100,101,102 | Stable clones from UTI89 Tn5 library ME-$P_{\sigma 70}$-mCherry- $P_{rhaB}$-*tse2*-kan-oriT-ME transformed with pRK24 | This study |
| Pooled recipient library for MAE VK8-93 | MG1655 ME-chlor-$P_{rhaB}$-*tse2*-ME | This study |
| Recipient single clones VK8-90-1 to VK8-90-948 | MG1655 ME-chlor-$P_{rhaB}$-*tse2*-ME | This study |
| Pooled recipient library with GFP plasmid for MAE<br>VK9-27 | MG1655 ME-chlor-$P_{rhaB}$-*tse2*-ME/pSLC-391 | This study |
| VK9-30 | Pooled transconjugant library of ~1400 cfu | This study |

| | | |
|---|---|---|
| VK9-30-1 to VK9-30-216 | 216 independent transconjugants | This study |
| VK10-85 | Transconjugant library of ~12000 cfu | This study |
| VK10-85-1 to VK10-85-168 | 168 independent transconjugants | This study |
| VK9-120-16<br>VK9-120-21<br>VK9-120-24<br>VK9-120-28<br>VK9-120-35<br>VK9-120-41<br>VK9-120-45<br>VK9-120-48<br>VK9-120-49<br>VK9-120-55<br>VK9-120-58<br>VK9-120-68<br>VK9-120-69<br>VK9-120-71<br>VK9-120-74<br>VK9-120-79<br>VK9-120-90<br>VK9-120-102 | Pod formers and non pod formers screened via PCR for different regions of the common 314kb locus transferred | This study |
| SLC-1222 | UTI89 UTI89 *yefM*::$P_{\sigma70}$-mCherry-$P_{rhaB}$-*tse2*-kan-oriT | This study |
| SLC-1223 | MG1655::chlor-$P_{rhaB}$-*tse2* downstream of *wcaM* | This study |
| SLC-1224 | SLC-1222/pRK24 | This study |
| VK10-90-1<br>VK10-90-2<br>VK10-90-5 to VK10-90-20<br>VK10-90-29<br>VK10-90-30<br>VK10-90-47 | $Serum^R$ transconjugants isolated from serum screen of VK10-85 transconjugant library | This study |
| VK10-90-63 | $Serum^S$ transconjugant isolated from serum screen of VK10-85 transconjugant library (-ve control) | This study |
| SLC-1235 | UTI89 UTI89 *yggX*::$P_{\sigma70}$- | |

| | | |
|---|---|---|
| | mCherry- $P_{rhaB}$-*tse2*-kan-oriT | |
| SLC-1236 | SLC-1235/pRK24 | |
| SLC-1237 | MG1655::chlor-$P_{rhaB}$-*tse2* downstream of *yqgA* | |
| SLC-1238 | P1 $Phage^{R}$ transconjugant | This study |
| SLC-1239 | P1 $Phage^{R}$ transconjugant | This study |
| VK9-53-5 | Pod forming and P1 phage resistant transconjugant | This study |
| VK9-53-6 | Pod forming transconjugant | This study |
| VK9-57-8 | Pod forming transconjugant | This study |
| VK9-57-10 | Pod forming transconjugant | This study |
| VK9-57-11 | Pod forming transconjugant | This study |
| VK10-26-4 | VK9-53-5 Δ*kpsFS*::kan | This study |
| VK10-27-2 | VK9-53-5 Δ*neuSD*::kan | This study |
| VK10-28-1 | VK9-53-5 Δ*kpsTM*::kan | This study |
| SLC-1246 | UTI89 Δ*chu* operon::FRT | This study |
| SLC-1244 | UTI89 Δ*auf* operon::kan | This study |
| VK9-88-3 | VK9-53-5 Δ*chu* operon::kan | This study |
| VK10-36-3 | VK9-57-11 Δ*chu* operon::FRT | This study |

**Table S2. List of plasmids**

| Plasmids | Genotype and/or relevant characteristics | Source/ reference |
|---|---|---|
| pACYC177 | | Rose RE, 1988 |
| pAH144 | CRIM plasmid with *aad* gene | Haldimann and Wanner, 2001 |

| | | |
|---|---|---|
| pAH120 | CRIM plasmid with $P_{rhaB}$ | Haldimann and Wanner, 2001 |
| pKD4 | | Datsenko and Wanner, 2000 |
| pKD46 | | Murphy and Campellone, 2003 |
| pKM208 | | Murphy and Campellone, 2003 |
| pSLC-130 (pXJ40_2) | mCherry with $P_{\sigma70}$ | Gift from Sohail Ahmed, Institute of Medical Biology (IMB), A*Star |
| pSLC-389 | *tse2* toxic gene cloned under $P_{rhaB}$ on pAH120 using NdeI and BamHI | This study |
| EZ-Tn5$^{TM}$ pMOD$^{TM}$-3 | EZ-Tn5$^{TM}$ pMOD$^{TM}$-3 <R6Kγori/MCS> transposon construction vectors | Epicentre - catalog number. TNP10623 |
| pSLC-294 | pBAD33 with $P_{\sigma70}$-vsfGFP-9 | Lab stock |
| pSLC-390 | Ez-Tn5 pMOD-3 <$P_{\sigma70}$-mCherry- $P_{rhaB}$-*tse2*-kan-oriT> | This study |
| pSLC-281 | chlor-$P_{rhaB}$-*tse2* | This study |
| pSLC-217 | kan-$P_{rhaB}$-*relE* | Khetrapal et al., 2015 |
| pRK24 | Bacterial strain: 51950 Conjugative helper plasmid | Addgene/ Natalie J Ma et al., 2014 |
| pSLC-391 | vsfGFP with Strep$^R$, Spec$^R$ and Chlor$^R$ | This study |
| pSLC-392 | pACYC177 with chu operon Kan$^R$ | This study |

**Table S3. List of primers**

| No. | Primer sequence | Notes |
|---|---|---|
| 1 | TGTGTAGGCTGGAGCTGCTTC | pKD4 universal primers |
| 2 | CATATGAATATCCTCCTTAG | |
| 3 | TTAACCAAATCGTCGTAAGCACACA GTAAATCACGCCAGGCCACCTCAAT GCTTGCAGTGGGCTTACATG | Fwd primer for amplification of kan-PrhaB-*relE* selection cassette from pSLC-217 for insertion at uti8+4887500 within the PAI II of UTI89 |
| 4 | GTTCTGAAGACGGAATCAGAAAAA TAATTCCAGAGTAAACCTGTCTGTT TTCAGAGCAGGATCGACGTCC | Rev primer for amplification of kan-PrhaB-*relE* selection cassette from pSLC-217 for insertion at uti8+4887500 within the PAI II of UTI89 |
| 5 | ACGTTTCACTCCGCGCCATA | Fwd test primer for insertion in PAI II |
| 6 | CGCATCATATGGGGGCACGT | Rev test primer for insertion in PAI II |
| 7 | AAGCCTGCAGCAAGCTGGCGAGTT CTCGATTGATCACCTGCTGGTTAAA ACTTCATTTAAATGGCGCGCC | Fwd primer for amplification of chlor-PrhaB-*tse2* selection cassette for insertion at K12+2113936 downstream of wcaM for selection of directed transfer of rfbUTI89 |
| 8 | GACGAATTGGCTCCGGTCGTCAAAC GCGCGCGCGAAAAAGTTGAACACG ACATATGAATATCCTCCTTAG | Rev primer for amplification of chlor-PrhaB-*tse2* selection cassette for insertion at K12+2113936 downstream of wcaM for selection of directed transfer of rfbUTI89 |
| 9 | TGGAGGAAGGCAAGCGCCGA | Fwd test primer for *WcaM* |
| 10 | ACAAATCCGGCTGGCTGGTG | Rev test primer for *WcaM* |
| 11 | CCTCTTTTGTACAGTTCATTGTACA ATGATGGGTGTTAATTAACTATTTA | Fwd primer for amplification of mCherry-PrhaB-*tse2*-kan-oriT selection cassette for |

| | | |
|---|---|---|
| | CTGTCTCTTATACACATCTCAACCA TCA | insertion at UTI8_+2228129 upstream of yefM for directed transfer of rfbUTI89 |
| 12 | AGTCTGATTTTTCGTCTCGCTGACA ATGTCTTGATCTACAAACTAATTAA AGCGCTTTTCCGCTGCA | Rev primer for amplification of mCherry-PrhaB-*tse2*-kan-oriT selection cassette for insertion at UTI8+2228129 upstream of *yefM* for directed transfer of rfbUTI89 |
| 13 | ATGATCTTCAACGGCTTTCA | Fwd test primer for *yefM* |
| 14 | TTTATGGCGATGGATTTGTT | Rev test primer for *yefM* |
| 15 | CTGTCTCTTATACACATCTCAACCA TCA | ME Plus 9 – 3′ primer for amplification of custom transposons on pMOD3 background |
| 16 | CTGTCTCTTATACACATCTCAACCC TGA | ME Plus 9 – 5′ primer for amplification of custom transposon on pMOD3 background |
| 17 | CTGTCTCTTATACACATCTCTTCATT TAAATGGCGCGCC | PO4-EZ-Tn5 pKD3 fwd for amplification of chlor-PrhaB-*tse2* for Tn5 mutagenesis |
| 18 | CTGTCTCTTATACACATCTGACATG GGAATTAGCCATGGTC | PO4-EZ-Tn5 pKD3 rev for amplification of chlor-PrhaB-*tse2* for Tn5 mutagenesis |
| 19 | GATCCTGGATCCACGATTGAACAAT ATAAAGAGGCCCAGCAACAGCCAG ACC | BamHI_UTI8+3912974_ChuS_fwd for amplification of Chu operon |
| 20 | GATCCTGTGCACGTGCGGTACTCCT GTTGGTTTTAATGATCGAGGACGAT CA | ApaLI_UTI8-3922002_hmuV_rev for amplification of Chu operon |
| 21 | TTATGCAGTAACGCCTTCCGGTATC AACGAAGCAATTTGCTTACGCCATT GTGTAGGCTGGAGCTGCTTC | Fwd primer for amplification of kan from pKD4 for knockout of chu operon |
| 22 | TTAATGATCGAGGACGATCAGTGGT GAATGGTTGGCAGGATCTTTAAGTA CATATGAATATCCTCCTTAG | Rev primer for amplification of kan from pKD4 for knockout of chu operon |
| 23 | GCAACAGCCAGACCTCCCGT | Fwd test primer for Chu operon knockout |
| 24 | TGGCAGGGGTGAAAGGCGTT | Rev test primer for Chu operon knockout |
| 25 | TTACCCCCTCGCCCGCTGATCGCGT GTAAAGGGTTGATAGACAAAACGA | Fwd primer for amplification of kan from pKD4 for knockout of auf operon |

| | | |
|---|---|---|
| | GGTGTAGGCTGGAGCTGCTTC | |
| 26 | ATGAAATTCAATTTATCTAATTTAT CCGCAGTATTACTGGCATCAGGTAT CATATGAATATCCTCCTTAG | Rev primer for amplification of kan from pKD4 for knockout of auf operon |
| 27 | ACAAATTGCTTCGGCGCCGG | Fwd test primer for auf operon knockout |
| 28 | GCCCCTCTTACAACTCCACG | Rev test primer for auf operon knockout |
| | TTAATATATATTATGTTGGCAGTTT GTTGTGTTCCCCCCATAATAAACCC CATATGAATATCCTCCTTAG | Rev primer for amplification of kan from pKD4 for knockout of *kpsST* |
| 29 | GTGACTGTGGCATTATTTCCGTGCA AAGGAGCTGATATGTCTGAAAGAC AGTGTAGGCTGGAGCTGCTTC | Fwd primer for amplification of kan from pKD4 for knockout of *kpsFS* |
| 30 | CTTTGAAGAGGATGGAAATG | Rev test primer for *kpsST* operon knockout |
| 31 | TGGTTCCCCTTCTCGCCTGT | Fwd test primer for *kpsST* operon knockout |
| 32 | TTACTCCCCCAAGAAAATCCTTTTA TCGTGCAAAGAGGGAGATGTTATAT GTGTAGGCTGGAGCTGCTTC | Fwd primer for amplification of kan from pKD4 for knockout of *neuSD* |
| 33 | ATGAGTAAAAAGTTAATAATATTTG GTGCGGGTGGTTTTTCAAAATCTAT CATATGAATATCCTCCTTAG | Rev primer for amplification of kan from pKD4 for knockout of *neuSD* |
| 34 | CCATCCTCTTCAAAGAAAAG | Fwd test primer for *neuSD* knockout |
| 35 | ACCTATAGTGGTTACATTCC | Rev test primer for *neuSD* knockout |
| 36 | CTATAGGTCTTTTTTGTAATCAGCA ATGGCTTCCGTAACATTTTTATAAA GTGTAGGCTGGAGCTGCTTC | Fwd primer for amplification of kan from pKD4 for knockout of *kpsTM* |
| 37 | ATGGCAAGAAGTGGATTTGAAGTTC AGAAAGTCACCGTAGAGGCATTATT CATATGAATATCCTCCTTAG | Rev primer for amplification of kan from pKD4 for knockout of *kpsTM* |

| | | |
|---|---|---|
| 38 | CCACCCGCACCAAATATTAT | Fwd test primer for *kpsTM* knockout |
| 39 | GCCAAGGGCGGTAGCGTACC | Rev test primer for *kpsTM* knockout |
| 40 | AGATTGTGCAGTCTGCAGTAAATT GAAGAAATTTGATTGACGAGACG AGGCTTCATTTAAATGGCGCGCC | Fwd primer for amplification of chlor-PrhaB-*tse2* selection cassette for insertion at K12+3108313 downstream of *yqgA* for selection of directed transfer of kps operon |
| 41 | GCTCTACCGACTGAGCTATCCGGG CAACGGGGCGCATTAAACCTGATT CGCATATGAATATCCTCCTTAG | Rev primer for amplification of chlor-PrhaB-*tse2* selection cassette for insertion at K12+3108313 downstream of *yqgA* for selection of directed transfer of kps operon |
| 42 | TCAACATGCTTCCAGCACTC | Fwd test primer for *yqgA* |
| 43 | ACTCTCGTTAACGTGGCGAG | Rev test primer for *yqgA* |
| 44 | TACGCACTGGCGCGCCGGTTTAGC GCGTGAGTCGATAAAGAGGATGA TTTCTGTCTCTTATACACATCTCAA CCATCA | Fwd primer for amplification of mCherry-PrhaB-tse2-kan-oriT selection cassette for insertion at UTI8+3281599 upstream of *yggX* for directed transfer of kps operon |
| 45 | TGACCTTCTGCTTCACGTTGCAGG AAAGTACAAAAAATCGTTCTGCTC ATAGCGCTTTTCCGCTGCA | Rev primer for amplification of mCherry-PrhaB-tse2-kan-oriT selection cassette for insertion at UTI8+3281599 upstream of *yggX* for directed transfer of kps operon |
| 46 | GCCGGTTTAGCGCGTGAGTC | Fwd test primer for *yggX* |
| 47 | GCTTGCGGTGCTCGGCATTC | Test test primer for *yggX* |
| 48 | AATGATACGGCGACCACCGAGATC TACACTCTTTCCCTACACGACGCTC TTCCGATCT ACTTGA acaagagcttcagggttg AGATGTG | TruSeq_Tn5pMOD3_AD008 TraDIS for UTI89 donor library |
| 49 | AATGATACGGCGACCACCGAGATC TACACTCTTTCCCTACACGACGCTC TTCCGATCTCAGATCTAAGGAGGAT ATTCATATGGACCAT | TruSeq_Tn5Cm_AD007 TraDIS primer for VK8-93-library |
| 50 | AATGATACGGCGACCACCGAGATC TACACTCTTTCCCTACACGACGCTC TTCCGATCTGATCAGTAAGGAGGAT ATTCATATGGACCAT | TruSeq_Tn5Cm_AD009 TraDIS primer for VK9-27-library |

| | | |
|---|---|---|
| 51 | AATGATACGGCGACCACCGAGATC TACACTCTTTCCCTACACGACGCTC TTCCGATCTATCACGTAAGGAGGAT ATTCATATGGACCAT | TruSeq_Tn5Cm_AD001 VK5-102-library |
| 52 | AATGATACGGCGACCACCGAGATC TACACTCTTTCCCTACACGACGCTC TTCCGATCTTTAGGCTAAGGAGGAT ATTCATATGGACCAT | TruSeq_Tn5Cm_AD003 1000 pooled colonies |
| 53 | AATGATACGGCGACCACCGAGATC TACACTCTTTCCCTACACGACGCTC TTCCGATCTTGACCATAAGGAGGAT ATTCATATGGACCAT | TruSeq_Tn5Cm_AD004 stable |
| 54 | AATGATACGGCGACCACCGAGATC TACACTCTTTCCCTACACGACGCTC TTCCGATCTACAGTGTAAGGAGGAT ATTCATATGGACCAT | TruSeq_Tn5Cm_AD005 unstable |
| 55 | CAAGCAGAAGACGGCATACGAGAT CGTGATGTGACTGGAGTTC | PCR primer index 1 for TraDIS |
